# The scRNA-seq expression profiling of the receptor ACE2 and the cellular protease TMPRSS2 reveals human organs susceptible to COVID-19 infection

**DOI:** 10.1101/2020.04.16.045690

**Authors:** Jing Qi, Yang Zhou, Jiao Hua, Liying Zhang, Jialin Bian, Beibei Liu, Zicen Zhao, Shuilin Jin

## Abstract

**Background:** COVID-19 caused by SARA-CoV-2 is a disaster sweeping over 200 countries, and more than 2,150,000 people are suffering from the disease and 140,000 people died. ACE2 is a receptor protein of SARS-CoV-2, and TMPRSS2 promotes virus proliferation and transmission. Some patients developed multiple organ dysfunction syndromes other than lungs. Therefore, studying the viral susceptibility of other organs is important for a deeper understanding of viral pathogenesis.

**Methods:** The advantage of scRNA-seq data is the identification of cell types by clustering the gene expression of cells. ACE2 and TMPRSS2 are highly expressed in AT2 of lungs, we compared the ACE2 and TMPRSS2 expression levels of cell types from 31 organs, with AT2 of lungs to evaluate the risk of the viral infection using scRNA-seq data.

**Findings:** For the first time, we found the brain, gall bladder, and fallopian tube are vulnerable to COVID-19 infection. Besides, the nose, heart, small intestine, large intestine, esophagus, testis and kidney are also identified to be high-risk organs with high expression levels of ACE2 and TMPRSS2. Moreover, the susceptible organs are grouped into three risk levels based on the TMPRSS2 expression. As a result, the respiratory system, digestive system and reproductive system are at the top-risk level to COVID-19 infection.

**Interpretation:** This study provides evidence for COVID-19 infection in the human nervous system, digestive system, reproductive system, respiratory system, circulatory system and urinary system using scRNA-seq data, which helps for the clinical diagnosis and treatment of patients.

## Introduction

In December 2019, a novel pneumonia coronavirus, recently named Coronavirus Disease 2019 (COVID-19) by World Health Organization (WHO), was first detected in several patients in Wuhan, China. Pneumonia spread widely in more than 200 countries, and more than 2,150,000 people were suffering from the disease and over 140,000 people died, posed a major threat to the global public health. COVID-19 is caused by the severe acute respiratory syndrome coronavirus 2 (SARS-CoV-2),^1^ which seriously damaged the respiratory system. Some patients developed acute respiratory infection symptoms, and even acute respiratory distress syndrome (ARDS), acute respiratory failure and other complications.^2,3^ ^2^On the other side, multiple organ dysfunction syndromes occurred in some patients and even led to death, suggesting that the virus invades other organs at the same time.^22^ SARS-CoV-2 enters the cell via angiotensin-converting enzyme II (ACE2), the receptor protein of SARS-CoV and NL63,^4–9^ while the cellular protease TMPRSS2 promotes the transmission during the viral infection.^8–12^ It is reasonable to predict the risk of organs vulnerable to COVID-19 infection using the expression level of ACE2 and TMPRSS2.

The advantage of scRNA-seq over bulk RNA sequence is the identification of cell types by clustering the gene expression of cells. To explore the potentially susceptible organs, previous studies analyzed the ACE2 expression of some human organs using scRNA-seq data. The gross anatomical observation indicates that the lesions caused by the novel coronavirus are mainly in the lung.^13^ ACE2 is mainly expressed in type II alveolar cells (AT2) in the lung,^14^ which implies AT2 cell is the cell type vulnerable to COVID-19 infection. Many researchers have obtained the susceptibility of other organs from different system using ACE2 expression in AT2 cells as a reference. Following this way, many respiratory organs were considered, and ACE2 was reported to be highly expressed in the nasal tissue, mouth, airway, and lung.^15–17^ The esophagus, large intestine (ileum and colon), and pancreas were identified as high-risk organs in the digestive system by the following papers.^16–19^ The kidney and bladder as major organs of urinary system were also indicated to be high ACE2-expressed.^17,20,21^ Besides, the testes and uterus were manifested to be susceptible organs, implying that the reproductive system was a potential route of viral infection.^22,23^

In this paper, the expression level of ACE2 and TMPRSS2 in different cell clusters of organs from nine major systems (including the respiratory system, digestive system, nervous system, endocrine system, reproductive system, circulatory system, urinary system, and motor systems) were obtained using the scRNA-seq data. For the first time, we found the brain (substantia nigra and cortex), fallopian tube, and gall bladder are vulnerable to COVID-19 infection; Besides, the lung, nose (nasal brushing epithelial cells, nasal turbinate epithelial cells, and nasal airway epithelial cells), heart, small intestine (jejunum, ileum, and duodenum), large intestine (rectum and colon), esophagus, testes, and kidney are predicted as high-risk organs under a more rigorous standard; Moreover, as the spike (S) protein initiated by TMPRSS2 is essential for the entry of the virus into the target cells and the transmission of the virus in the infected host, we used the expression level of TMPRSS2 to predict the risk level of each susceptible organ, and found that the respiratory system, digestive system, and reproductive system are the top level vulnerable to COVID-19 infection.

## Methods

The scRNA-seq data of healthy human available are collected for the analysis, including 31 organs from nine major human systems (Table 1), and the details of the data resources are in Supplementary File Table S1.

**Table 1:**
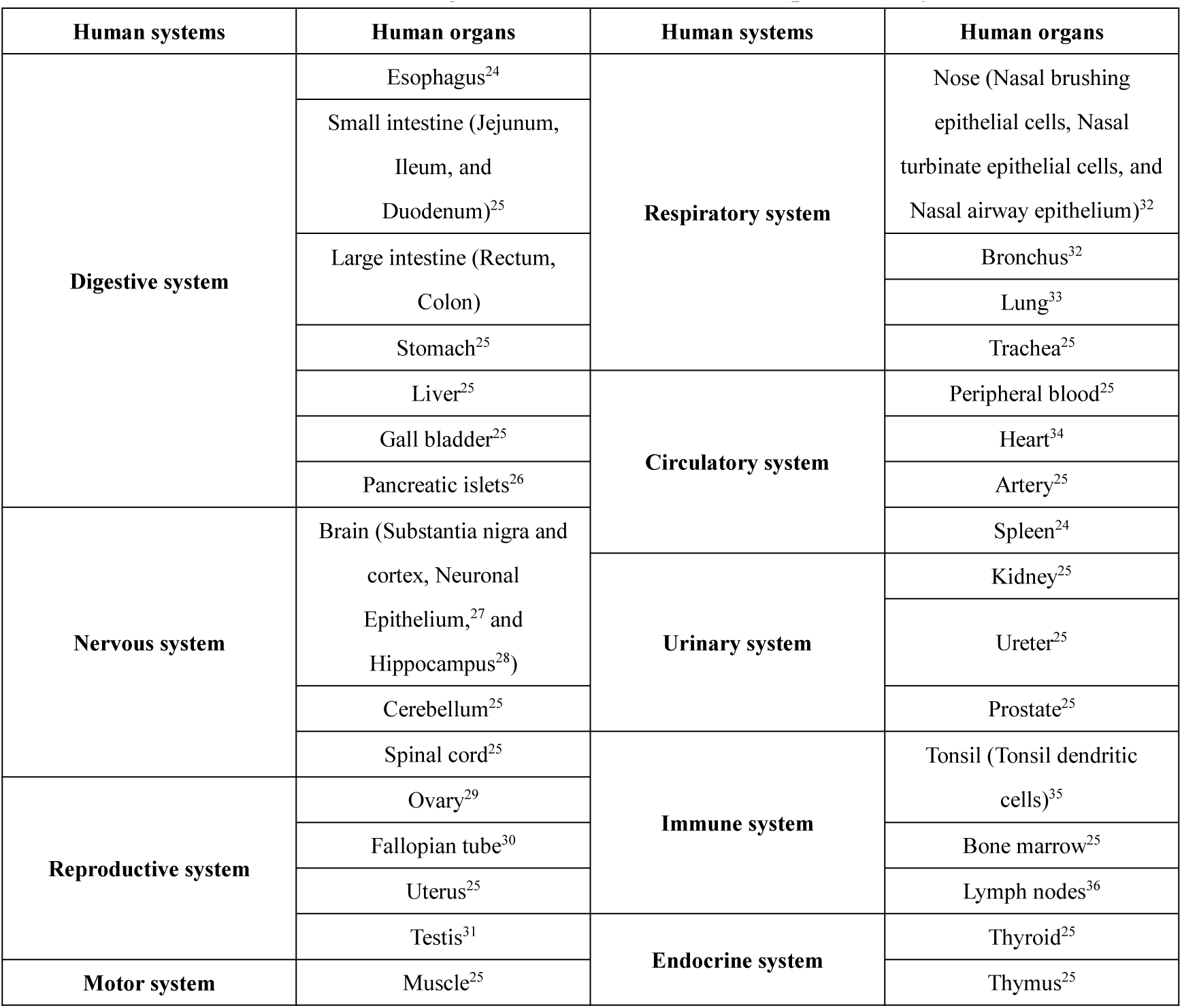
Human organs available for scRNA-seq data analysis

R package, Seurat, was applied to analyze the scRNA-seq data (details see Supplementary File Text S1). The expression level of ACE2 in AT2 cells is the primary reference to infer the susceptibility of other organs and tissues. The ratio of ACE2-expressed cells over total AT2 cells is 0•79% which is the mean of 8 samples. To determine the susceptibility of the organs, the expression level of TMPRSS2 is another reference. The cell cluster, in which TMPRSS2 is expressed and ACE2-expressed cell ratio is equal to or greater than AT2 cells at the same time, is identified as highly susceptible cells, and the corresponding organs are vulnerable to COVID-19.

As the TMPRSS2 is a key protease to help the SARS-CoV-2 enter the target cells, and the expression level of TMPRSS2 implies the sensitivity of the cell to COVID-19 infection. Therefore, the susceptible organs were sorted into three groups by the ratios of TMPRSS2-expressed cells. In details, the cells with TMPRSS2-expressed ratio greater 20% are defined as level 1, which is at the highest risk; the cells with ratio more than 5% and less than 20% are level 2, which means a higher risk; the rest are level 3, an existing risk of infection.

## Results

### Respiratory System

The scRNA-seq data of the lung, nose, trachea, and bronchus in the respiratory system were collected for the analysis. In the lung, AT2 cells contain an average of 0.79% ACE2-expressed cells across 8 samples (Fig. 1a, Fig. 1b, Supplementary File: Figure S1-S8, Table 2), and the expression levels of ACE2 and TMPRSS2 are high in AT2 cells (Fig. 1c, Fig. 1d). The data of nose (nasal brushing epithelial cells, nasal turbinate epithelial cells, and nasal airway epithelial cells) contains ACE2-expressed and TMPRSS2-expressed cell clusters (Supplementary File: Figure S9-S11), and the ratios of ACE2-expressed cells of these cell clusters are all over 0.79% (Table 2), thus the nose is identified as the high-risk organ. Low ratio of ACE2-expressed cells in the bronchus and trachea means they are low-risk infection organs (Supplementary File: Figures S12-S13).

**Table 2:** Cell types with high expression levels of ACE2 and TMPRSS2 in respiratory system

| Organs | Cell type | Ratio of ACE2-expressed cells | Ratio of TMPRSS2-expressed cells |
| --- | --- | --- | --- |
| Lung | AT2 cells | 0.79% | 21.2% |
| Nose (Nasal turbinate epithelial cells) | Mesenchymal stromal cells | 1.7% | 8.5% |
|  | Plasma cells | 2.0% | 8.8% |
| Nose (Nasal brushing epithelial cells) | Plasma cells | 4% | 13.8% |
| Nose (Nasal airway epithelial cells) | Nasal airway epithelial cells | 8.4% | 23.1% |

**Fig. 1.**
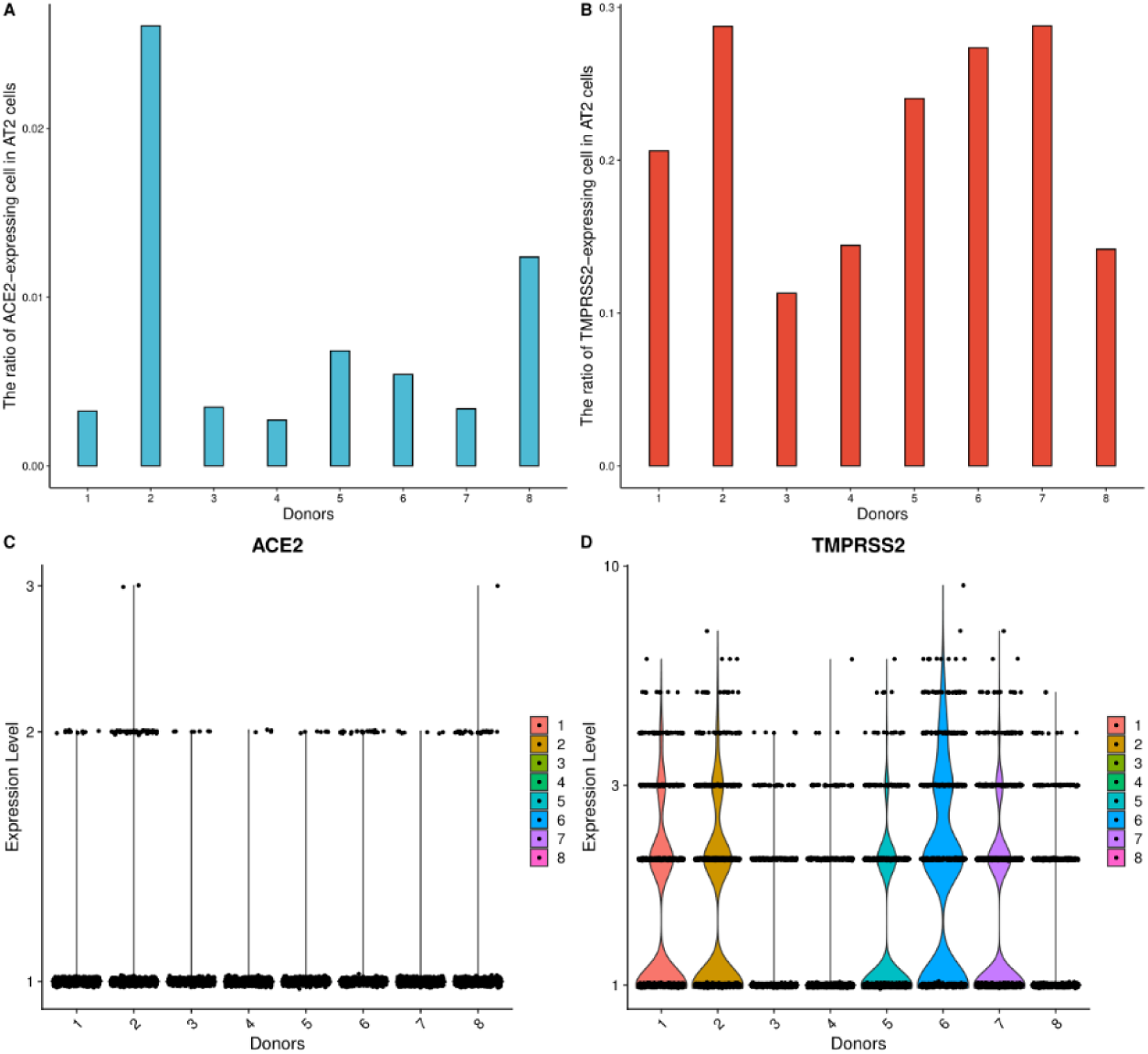
High ACE2 and TMPRSS2 expression level of AT2 cells in the lung. **a**) The ratio of ACE2-expressed cells in AT2 cells from 8 samples. **b**) The ratio of TMPRSS2-expressed cells in AT2 cells from 8 samples. **c**) The expression distribution of ACE2 in AT2 cells across 8 samples. **d**) The expression distribution of TMPRSS2 in AT2 cells across 8 samples.

### Digestive system

The scRNA-seq data of jejunum, ileum, duodenum, rectum, colon, esophagus, gall bladder, pancreatic islets, liver, and stomach from the digestive system were collected for the analysis. The primordium cells from the gall bladder contain 2.6% TMPRSS2-expressed cells and 2.2% ACE2-expressed cells (Fig. 2, Table 3), which means the gall bladder is vulnerable to the COVID-19 infection. Moreover, small intestine (jejunum, ileum, and duodenum), large intestine (rectum and colon), and esophagus are identified as high-risk organs (Supplementary File: Figures S14–S19, Table 3). However, no cell clusters from the liver, stomach, and pancreatic islets data show high ACE2 and TMPRSS2 expression level (Supplementary File: Figures S20-S22), which demonstrates a low infection risk.

**Table 3:** Cell types with high expression levels of ACE2 and TMPRSS2 in digestive system

| Organs | Cell type | Ratio of ACE2-expressed cells | Ratio of TMPRSS2-expressed cells |
| --- | --- | --- | --- |
| **Gall bladder** | Primordium cell | 2.2% | 2.6% |
| **Small intestine (Jejunum)** | Enterocyte progenitor cell | 14.0% | 6.6% |
|  | Goblet cell | 9.1% | 2.6% |
| **Small intestine (Ileum)** | Intestinal epithelial stem cell | 1.5% | 4.2% |
|  | Enterocyte progenitor cell | 2.4% | 3.6% |
| **Small intestine (Duodenum)** | LGR5+ stem cell | 5.2% | 4.5% |
|  | Intestinal epithelial stem cell | 3.9% | 5.8% |
|  | Enterocyte progenitor cell | 4.4% | 6.8% |
|  | Tuft progenitor cell | 7.4% | 6.6% |
|  | Enteroendocrine cell | 9.4% | 3.8% |
| **Large intestine (Rectum)** | Goblet progenitor cell | 2.8% | 62.5% |
|  | MKI67+ progenitor cell | 10.3% | 75.4% |
|  | Enterocyte | 13.2% | 77.5% |
|  | Goblet cell | 18.2% | 87.3% |
| **Large intestine (Colon)** | Enterocyte | 5.7% | 52.1% |
|  | Goblet cell | 6.7% | 52.4% |
| **Esophagus** | Secretory progenitor cell | 1.5% | 5.0% |

**Fig. 2.**
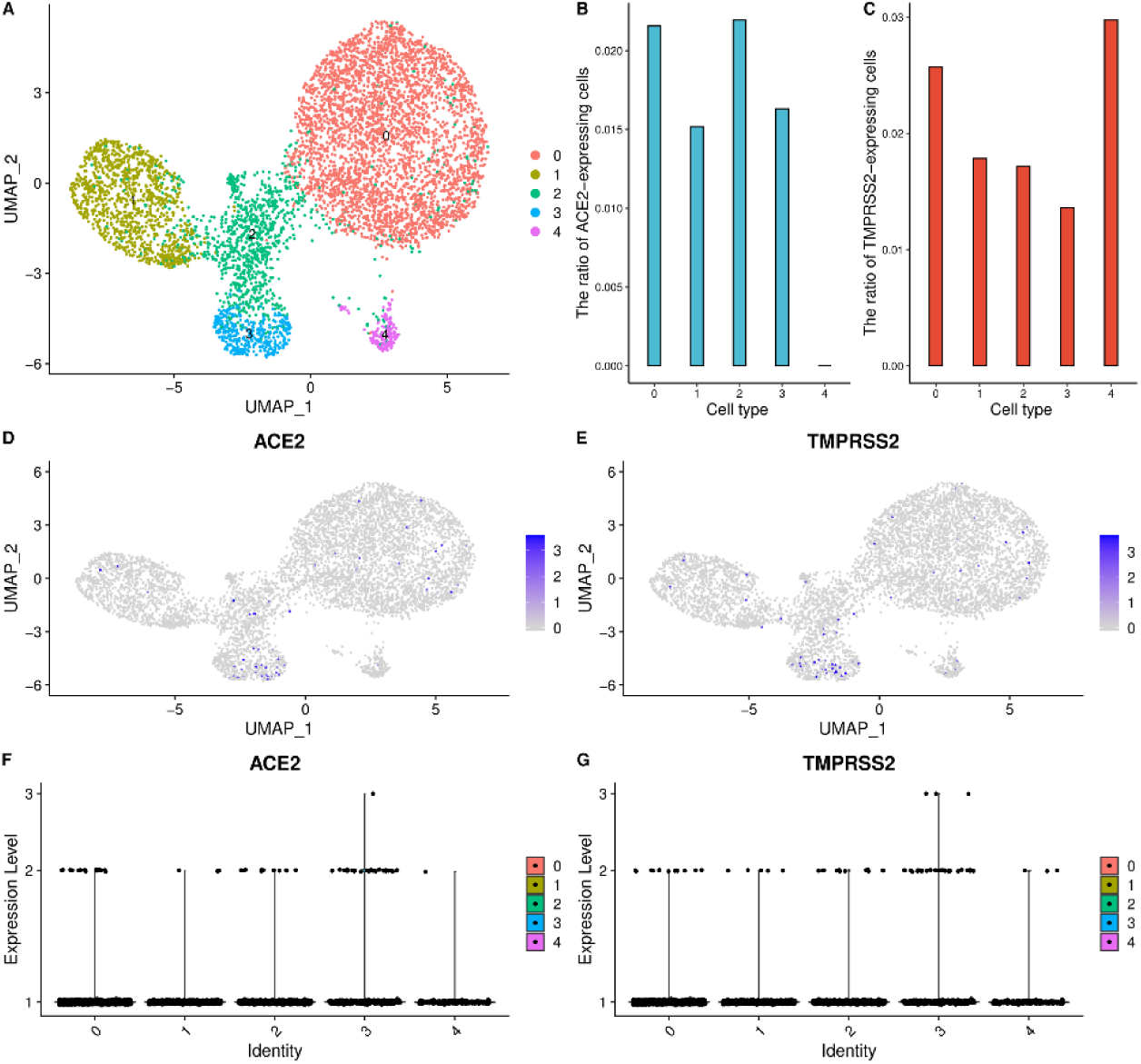
High ACE2 and TMPRSS2 expression level of the primordium cells in the gall bladder. **a**) UMAP visualization of clustering results for gall bladder cells. **b**) The ratio of ACE2-expressed cells in each cell cluster. **c**) The ratio of TMPRSS2-expressed cells in each cell cluster. **d**) ACE2 expression level in each cell cluster on the UMPA plot. **e**) TMPRSS2 expression level in each cell cluster on the UMPA plot. **f**) The expression distribution of ACE2 across each cell cluster. **g**) The expression distribution of TMPRSS2 across each cell cluster.

### Nervous System

The scRNA-seq of substantia nigra and cortex, hippocampus, cerebellum, spinal cord, and neuronal epithelium from the nervous system were collected to infer the susceptibility of the organs. The analysis results show that ACE2 is expressed in the oligodendrocyte precursor cells and astrocytes of the substantia nigra and cortex with a high level, and TMPRSS2 is expressed as well. In details, astrocytes contain 1.9% ACE2-expressed cells and oligodendrocyte precursor cells contain 1.6% ACE2-expressed cells (Fig. 3, Table 4). Therefore, the substantia nigra and cortex are predicted as high-risk tissues, and the brain is identified as high-risk organs. For the analysis of other organs, cells from the hippocampus have low expression level of TMPRSS2 and ACE2, and the cerebellum, spinal cord, and neuronal epithelium data show zero expression of TMPRSS2 and ACE2 (Supplementary File: Figures S23-S26), which demonstrates a low infection risk of these organs.

**Table 4:**
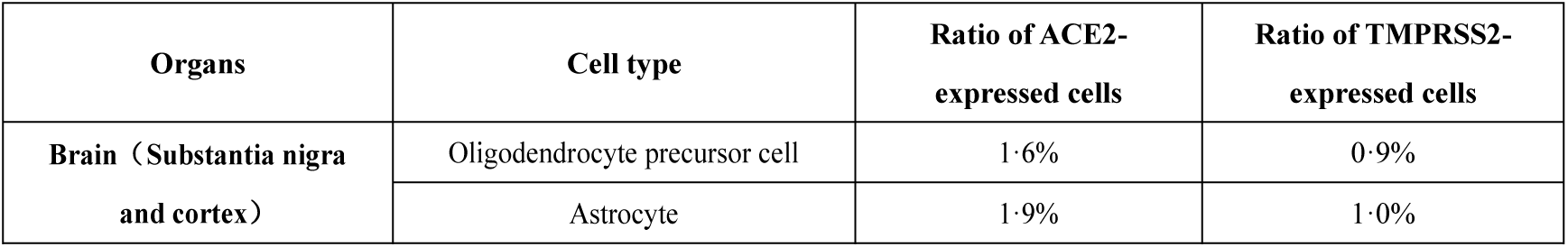
Cell types with high expression levels of ACE2 and TMPRSS2 in nervous system

**Fig. 3.**
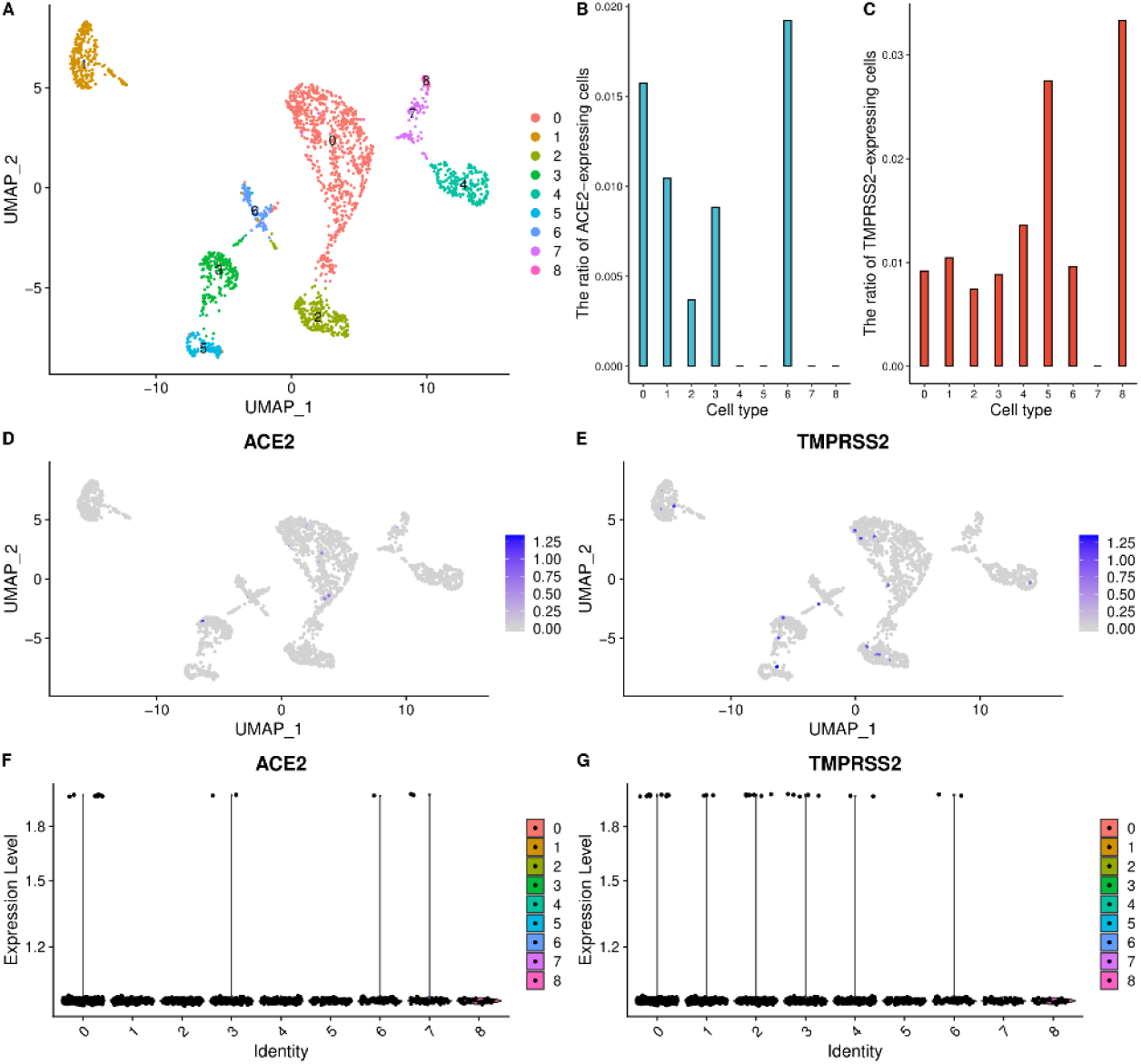
High ACE2 and TMPRSS2 expression level of oligodendrocyte precursor cells and astrocytes in the substantia nigra and cortex. **a**) UMAP visualization of clustering results for the substantia nigra and cortex cells. **b**) The ratio of ACE2-expressed cells in each cell cluster. **c**) The ratio of TMPRSS2-expressed cells in each cell cluster. **d**) ACE2 expression level in each cell cluster on the UMPA plot. **e**) TMPRSS2 expression level in each cell cluster on the UMPA plot. **f**) The expression distribution of ACE2 across each cell cluster. **g**) The expression distribution of TMPRSS2 across each cell cluster.

### Reproductive system

The scRNA-seq data of the testis, fallopian tube, ovary, and uterus from the reproductive system were collected for analysis. The ratios of the TMPRSS2-expressed cell and the ACE2-expressed cell in the epithelial cells of the fallopian tube are 26.5% and 1.4% respectively (Fig. 4, Table 5), and the ovarian somatic cells contain 1% TMPRSS2-expressed cells and 1% ACE2-expressed cells, so the fallopian tube is identified as a high-risk organ. The testis is also identified as a high-risk organ because of the high expression level of TMPRSS2 and ACE2 (Supplementary File: Figure S27, Table 5). Low ratios of ACE2- expressed cells in the ovary and uterus mean the ovary and uterus are low infection risk organs (Supplementary File: Figures S28-S29).

**Table 5:**
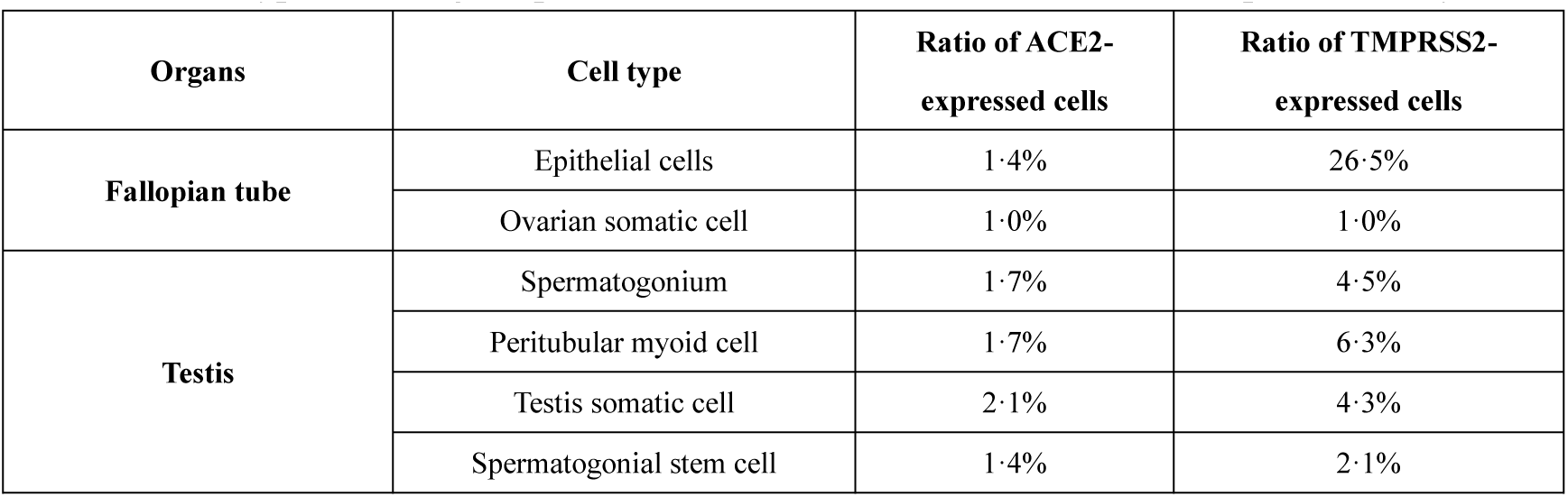
Cell types with high expression levels of ACE2 and TMPRSS2 in reproductive system

**Fig. 4.**
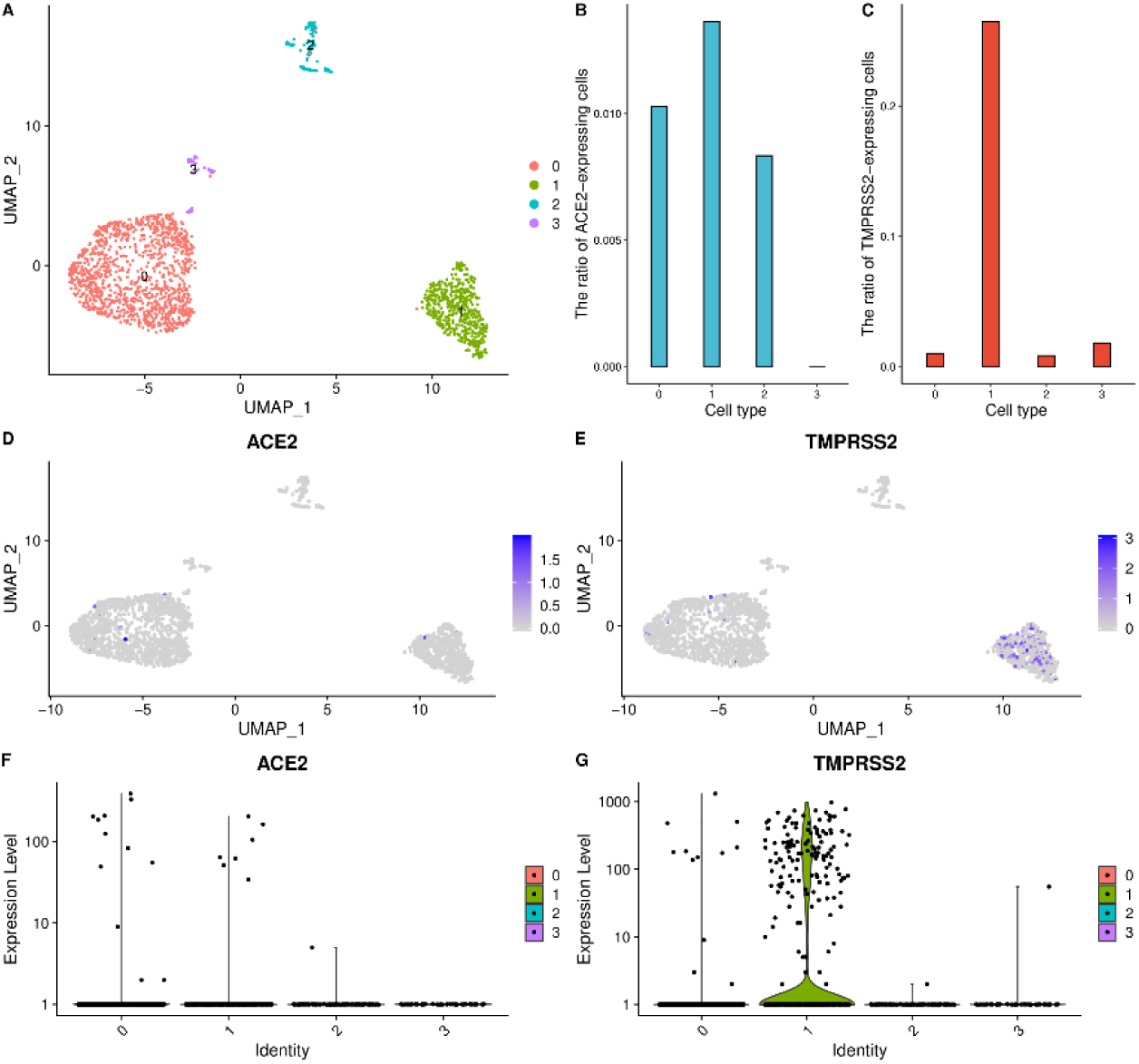
High ACE2 and TMPRSS2 expression level of epithelial cells and ovarian somatic cells in the fallopian tube. **a**) UMAP visualization of clustering results for the fallopian tube cells. **b**) The ratio of ACE2-expressed cells in each cell cluster. **c**) The ratio of TMPRSS2-expressed cells in each cell cluster. **d**) ACE2 expression level in each cell cluster on the UMPA plot. **e**) TMPRSS2 expression level in each cell cluster on the UMPA plot. **f**) The expression distribution of ACE2 across each cell cluster. **g**) The expression distribution of TMPRSS2 across each cell cluster.

### Circulatory system

The data of the heart, spleen, and artery from the circulatory system were collected to infer the susceptibility of these organs. The cardiomyocytes and cardiovascular progenitor cells from heart contain 6.6% and 12.5% ACE2-expressed cells respectively, and the TMPRSS2 is expressed in both cell clusters as well. Consequently, the heart is considered as a high-risk organ (Supplementary File: Figure S30, Table 6). Nevertheless, almost no cells of the spleen, artery, and peripheral blood data show high TMPRSS2 and ACE2 expression levels, which means they are low-risk organs (Supplementary File: Figures S31-S33).

**Table 6:** Cell types with high expression levels of ACE2 and TMPRSS2 in circulatory system

| Organs | Cell type | Ratio of ACE2-expressed cells | Ratio of TMPRSS2-expressed cells |
| --- | --- | --- | --- |
| Heart | Cardiomyocyte | 6.6% | 0.8% |
|  | Cardiovascular progenitor cell | 12.5% | 0.4% |

### Urinary System

The scRNA-seq data of the kidney, ureter, and prostate from the urinary system were utilized for the data analysis. The analysis results of the kidney scRNA-seq data show high ACE2 and TMPRSS2 expression levels in the nephron epithelial cells, epithelial cells, endothelial cells, and mesangial cells. Particularly, the ratios of TMPRSS2-expressed are 10.7%, 9.6%, 12.8%, and 14.5% respectively, and the ratios of ACE2- expressed are 2.7%, 2.7%, 2.7%, and 3.0% respectively (Supplementary File: Figure S34, Table 7). Therefore, the kidney is considered as high-risk organs. In addition, the ACE2 is not expressed in the ureter and prostate cells (Supplementary File: Figures S35-S36), and they are predicted as low-risk infection organs.

**Table 7:**
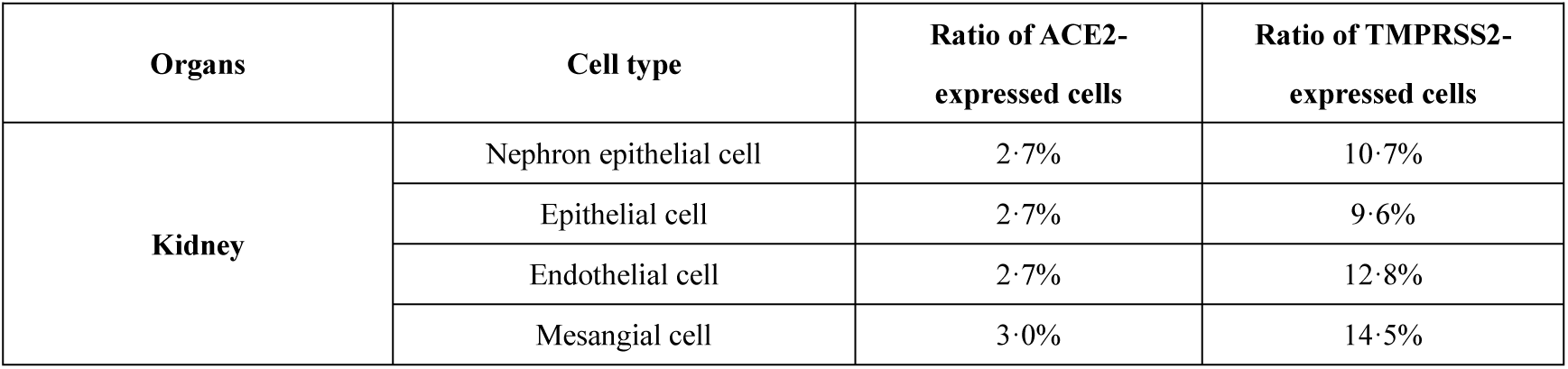
Cell types with high expression levels of ACE2 and TMPRSS2 in urinary system

### Endocrine system, Motor system and Immune system

The susceptibility of organs from the endocrine system, immune system, and motor system was also concerned. In the endocrine system, almost no cells from the thyroid gland data show high ACE2 and TMPRSS2 expression levels, and the thymus gland data contains no ACE2-expressed cells (Supplementary File: Figures S37-S38). Hence, they are not high-risk organs. Likewise, few ACE2-expressed cells are found in the muscle from the motor system (Supplementary File: Figure S39). There is no ACE2 expression in the lymph nodes, tonsil (tonsil dendritic cells), and bone marrow data from the immune system (Supplementary File: Figures S40- S42), which mean they are low-risk organs.

### The risk levels of susceptible organs

Based on TMPRSS2 expression level, we grouped the susceptible organs into three risk levels. In table 8, it is shown that lung, large intestine (colon and rectum), fallopian tube, and nose (nasal airway epithelium) are the most susceptible organs with TMPRSS2-expressed ration over 20%, and the result indicates the SARS-CoV-2 mainly attacks the respiratory system, the digestive system, and the reproductive system. The kidney, Small intestine (duodenum and jejunum), and testis are susceptible organs with moderate risk. In addition, the esophagus, gall bladder, brain (substantia nigra and cortex), and heart are identified to be the potentially susceptible organs.

**Table 8:** Risk level of organs infection by SARS-CoV-2

| Organs | Systems | Ratios of TMPRSS2-expressed cells | Cell types | Risk levels |
| --- | --- | --- | --- | --- |
| **Lung** | Respiratory system | 0.212 | AT2 | 1 |
| **Large intestine (Rectum)** | Digestive system | 0.873 | Goblet cell | 1 |
| **Large intestine (Colon)** | Digestive system | 0.524 | Goblet cell | 1 |
| **Fallopian tube** | Reproductive system | 0.265 | Epithelial cell | 1 |
| **Nose (Nasal airway epithelium)** | Respiratory system | 0.234 | All cells | 1 |
| **Nose (Nasal brushing epithelial cells)** | Respiratory system | 0.188 | Plasma cell | 2 |
| **Kidney** | Urinary system | 0.145 | Mesangial cell | 2 |
| **Nose (Nasal turbinate epithelial cells)** | Respiratory system | 0.088 | Plasma cell | 2 |
| **Small intestine (Duodenum)** | Digestive system | 0.068 | Enterocyte progenitor cell | 2 |
| **Small intestine (Jejunum)** | Digestive system | 0.066 | Enterocyte progenitor cell | 2 |
| **Testis** | Reproductive system | 0.063 | Peritubular myoid cell | 2 |
| **Esophagus** | Digestive system | 0.049 | Secretory progenitor cell | 3 |
| **Small intestine (Ileum)** | Digestive system | 0.042 | Intestinal epithelial stem cell | 3 |
| **Gall bladder** | Digestive system | 0.026 | Primordium cell | 3 |
| **Brain (Substantia nigra and cortex)** | Nervous system | 0.010 | Astrocyte | 3 |
| **Heart** | Circulatory system | 0.008 | Cardiomyocyte | 3 |

## Discussion

The clinical symptoms of patients infected by COVID are mainly manifested in the respiratory system and digestive system, including cough, shortness of breath, dyspnea, and diarrhea. However, some patients also developed symptoms such as heart damage and kidney failure, indicating that the virus affected the normal function of the circulatory and urinary systems. We investigated the susceptibility of the organs and tissues in various human systems based on the scRNA-seq data analysis. In details, 31 organs from nine major human systems were considered, in which 11 organs are identified to be susceptible to the virus. Moreover, we classified these susceptible organs into three levels in terms of their risk, which provide novel ideas for the follow-up detection of virus, treatment, and the monitoring of recrudescence.

However, due to the limitation of the data, only partial organs or tissues were involved, so the susceptible organs may be missed since the mechanism of cells infected by the COVID-19 is still not completely known. Furthermore, these organs reported to be susceptibility by the scRNA-seq data, should be further confirmed by the clinical observation and biological experiments.

## Supporting information

Supplementary Table, Text, and Figrues

## Authors’ contributions

Shuilin Jin and Jing Qi conceived and designed the study. Jiao Hua and Liying Zhang performed the data analysis. Yang Zhou, Jianlin Bian, and Beibei Liu collected for the available data. Jing Qi and Yang Zhou wrote the paper. Shuilin Jin, Jing Qi, Yang Zhou, Jiao Hua, Liying Zhang, Jialin Bian, Beibei Liu, and Zicen Zhao revised the manuscript. All authors read and approved the final manuscript.

## Acknowledgments

This work was funded by the Natural Science Foundation of China (Grant No. 11971130).

