## Supplementary Table, Text, and Figrues for "The scRNA-seq expression profiling of the receptor ACE2 and the cellular protease TMPRSS2 reveals human organs susceptible to COVID-19 infection"

**Table S1 The data resources of organs and tissues**

| Human systems | Human tissues | Data resources |
|---|---|---|
| **Digestive system** | Esophagus | https://www.tissuestabilitycellatlas.org/ |
| | Small intestine(Jejunum) | GEO: GSE134355 |
| | Small intestine(Ileum) | GEO: GSE134355 |
| | Small intestine(Duodenum) | GEO: GSE134355 |
| | Large intestine(Rectum) | GEO: GSE125970 |
| | Large intestine(Colon) | GEO: GSE125970 |
| | Stomach | GEO: GSE134355 |
| | Liver | GEO: GSE134355 |
| | Gall bladder | GEO: GSE134355 |
| | Pancreatic islets | GEO: GSE114297 |
| **Nervous system** | Brain(Substantia nigra and cortex) | GEO: GSE140231 |
| | Brain (Neuronal epithelium) | GEO: GSE81475 |
| | Brain (Hippocampus) | GEO: GSE119212 |
| | Cerebellum | GEO: GSE134355 |
| | Spinal cord | GEO: GSE134355 |
| **Reproductive system** | Ovary | GEO: GSE118127 |
| | Fallopian tube | GEO: GSE139079 |
| | Uterus | GEO: GSE134355 |
| | Testis | GEO: GSE112013 |
| **Motor system** | Muscle | GEO: GSE134355 |
| **Respiratory system** | Nose (Nasal brushing epithelial cells) | GEO: GSE121600 |
| | Nose (Nasal turbinate epithelial cells) | GEO: GSE121600 |
| | Nose (nasal airway epithelium) | GEO: GSE103518 |
| | Bronchus | GEO: GSE121600 |
| | Lung | GEO: GSE122960 |
| | Trachea | GEO: GSE134355 |
| **Circulatory system** | Peripheral blood | GEO: GSE134355 |
| | Heart | GEO: GSE106118 |
| | Artery | GEO: GSE134355 |
| | Spleen | https://www.tissuestabilitycellatlas.org/ |
| **Urinary system** | Kidney | GEO: GSE134355 |
| | Ureter | GEO: GSE134355 |
| | Prostate | GEO: GSE134355 |
| **Immune system** | Tonsil(Tonsil dendritic cells) | GEO: GSE115006 |
| | Bone marrow | GEO: GSE134355 |
| | Lymph nodes | GEO: GSE124494 |
| **Endocrine system** | Thyroid | GEO: GSE134355 |
| | Thymus | GEO: GSE134355 |

### Text S1 The pipeline of data analysis

The R package, Seurat, was used to analyze the scRNA-seq datasets in the following three steps. Step 1, filter the low-quality cells, according to the number of expressed genes, counts and cells with mitochondrial content, and reserve the cells within the range of $\mu - \sigma$ and $\mu + \sigma$ (where $\mu$ is the mean and $\sigma$ is the standard deviation of the numbers). Step 2, normalize the datasets by logarithmic transformation and carry out the downstream analysis with the top 2,000 most variable genes. Besides, to avoid the influence of extremely high expressed genes, the downstream analysis was given the same weight for gene expression by scaling the data. Step 3, perform the principal component analysis (PCA) on count matrix and use the top 10 principal components for the cell clustering analysis, and then label the Subtypes of cells by the t-test between the differential expression genes and marker genes of the cells.

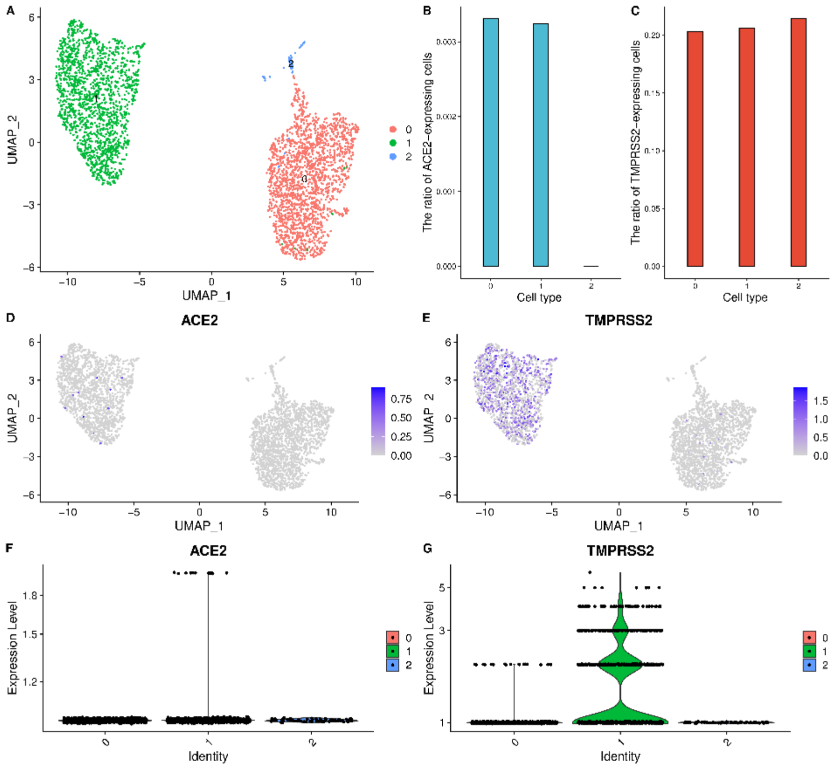


**Figure S1 The lung scRNA-seq data analysis results (donor 1). a**) UMAP visualization of clustering results for the lung cells. **b**) The ratio of ACE2-expressed cells in each cell cluster. **c**) The ratio of TMPRSS2-expressed cells in each cell cluster. **d**) ACE2 expression level in each cell cluster on the UMPA plot. **e**) TMPRSS2 expression level in each cell cluster on the UMPA plot. **f**) The expression distribution of ACE2 across each cell cluster. **g**) The expression distribution of TMPRSS2 across each cell cluster.

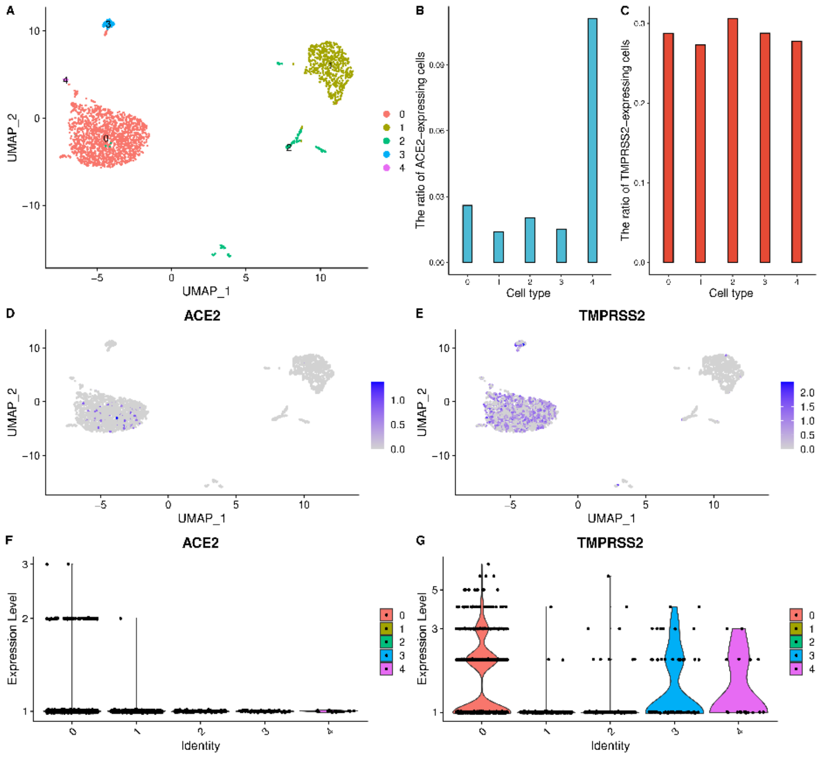


**Figure S2 The lung scRNA-seq data analysis results (donor 2). a**) UMAP visualization of clustering results for the lung cells. **b**) The ratio of ACE2-expressed cells in each cell cluster. **c**) The ratio of TMPRSS2-expressed cells in each cell cluster. **d**) ACE2 expression level in each cell cluster on the UMPA plot. **e**) TMPRSS2 expression level in each cell cluster on the UMPA plot. **f**) The expression distribution of ACE2 across each cell cluster. **g**) The expression distribution of TMPRSS2 across each cell cluster.

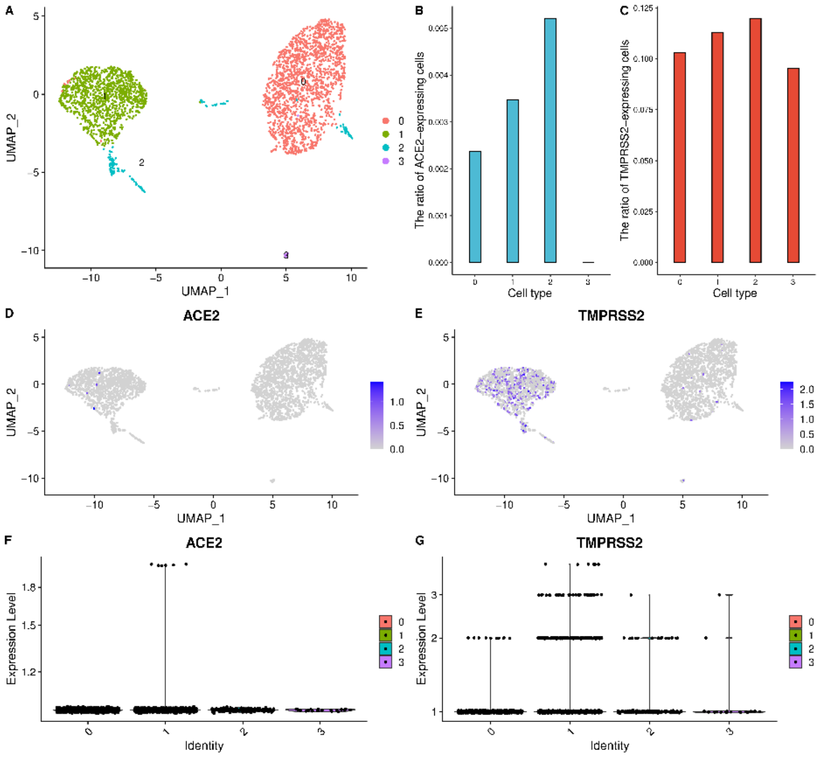


**Figure S3 The lung scRNA-seq data analysis results (donor 3). a**) UMAP visualization of clustering results for the lung cells. **b**) The ratio of ACE2-expressed cells in each cell cluster. **c**) The ratio of TMPRSS2-expressed cells in each cell cluster. **d**) ACE2 expression level in each cell cluster on the UMPA plot. **e**) TMPRSS2 expression level in each cell cluster on the UMPA plot. **f**) The expression distribution of ACE2 across each cell cluster. **g**) The expression distribution of TMPRSS2 across each cell cluster.

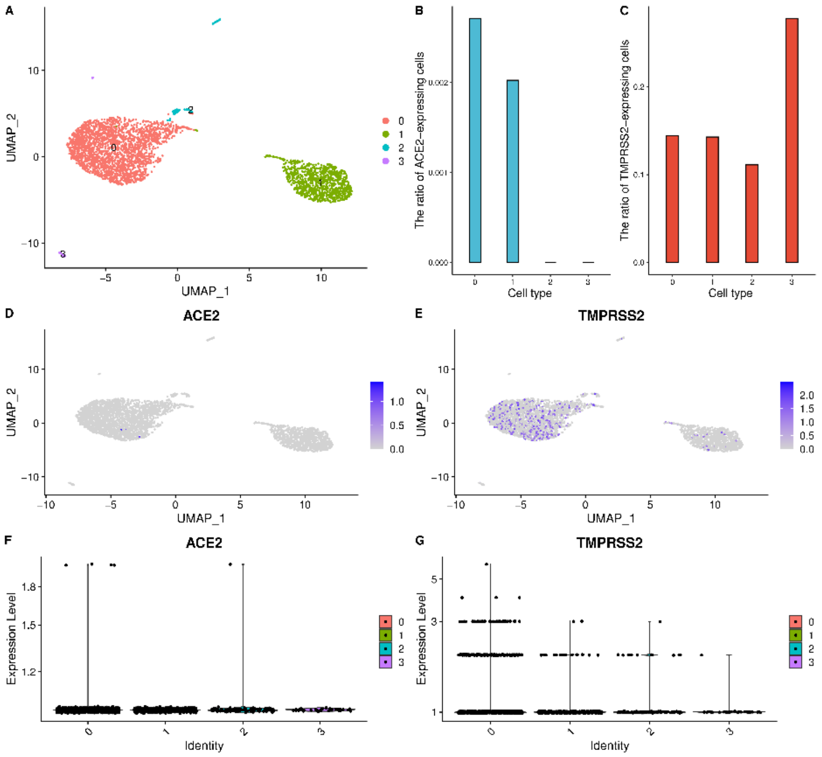


**Figure S4 The lung scRNA-seq data analysis results (donor 4). a**) UMAP visualization of clustering results for the lung cells. **b**) The ratio of ACE2-expressed cells in each cell cluster. **c**) The ratio of TMPRSS2-expressed cells in each cell cluster. **d**) ACE2 expression level in each cell cluster on the UMPA plot. **e**) TMPRSS2 expression level in each cell cluster on the UMPA plot. **f**) The expression distribution of ACE2 across each cell cluster. **g**) The expression distribution of TMPRSS2 across each cell cluster.

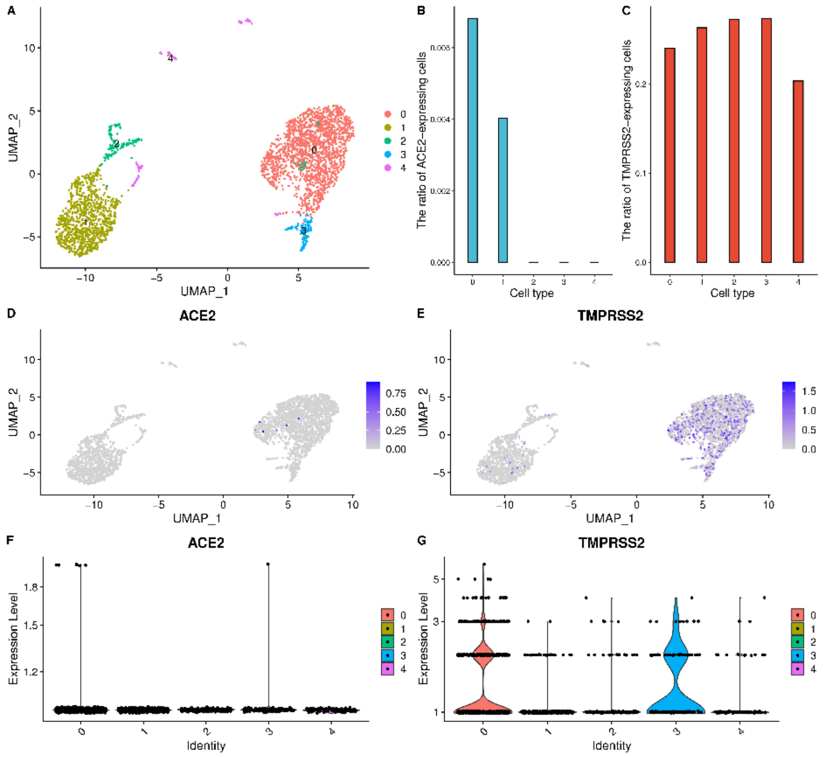


**Figure S5 The lung scRNA-seq data analysis results (donor 5). a**) UMAP visualization of clustering results for the lung cells. **b**) The ratio of ACE2-expressed cells in each cell cluster. **c**) The ratio of TMPRSS2-expressed cells in each cell cluster. **d**) ACE2 expression level in each cell cluster on the UMPA plot. **e**) TMPRSS2 expression level in each cell cluster on the UMPA plot. **f**) The expression distribution of ACE2 across each cell cluster. **g**) The expression distribution of TMPRSS2 across each cell cluster.

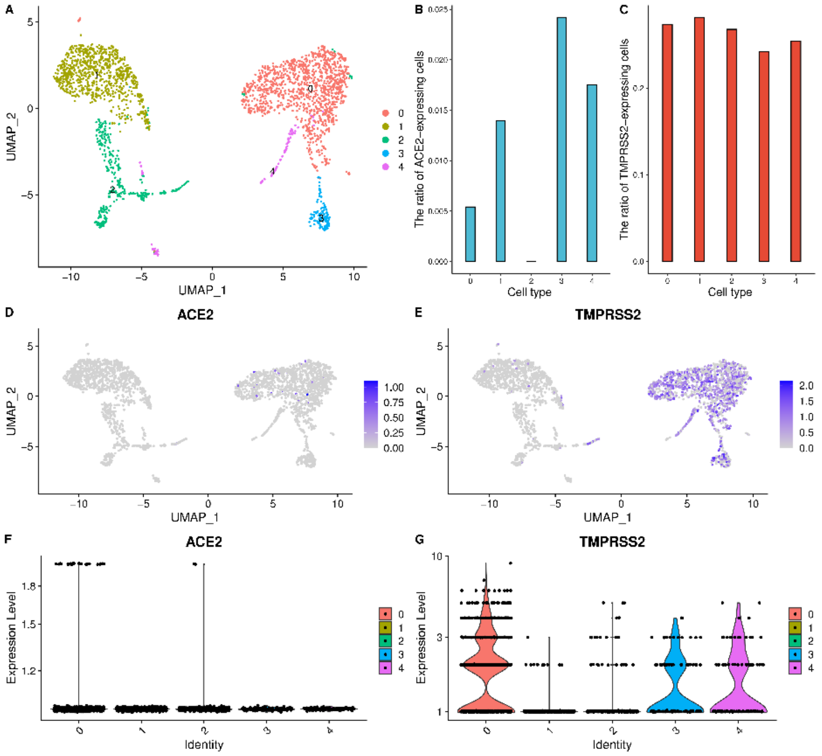


**Figure S6 The lung scRNA-seq data analysis results (donor 6). a**) UMAP visualization of clustering results for the lung cells. **b**) The ratio of ACE2-expressed cells in each cell cluster. **c**) The ratio of TMPRSS2-expressed cells in each cell cluster. **d**) ACE2 expression level in each cell cluster on the UMPA plot. **e**) TMPRSS2 expression level in each cell cluster on the UMPA plot. **f**) The expression distribution of ACE2 across each cell cluster. **g**) The expression distribution of TMPRSS2 across each cell cluster.

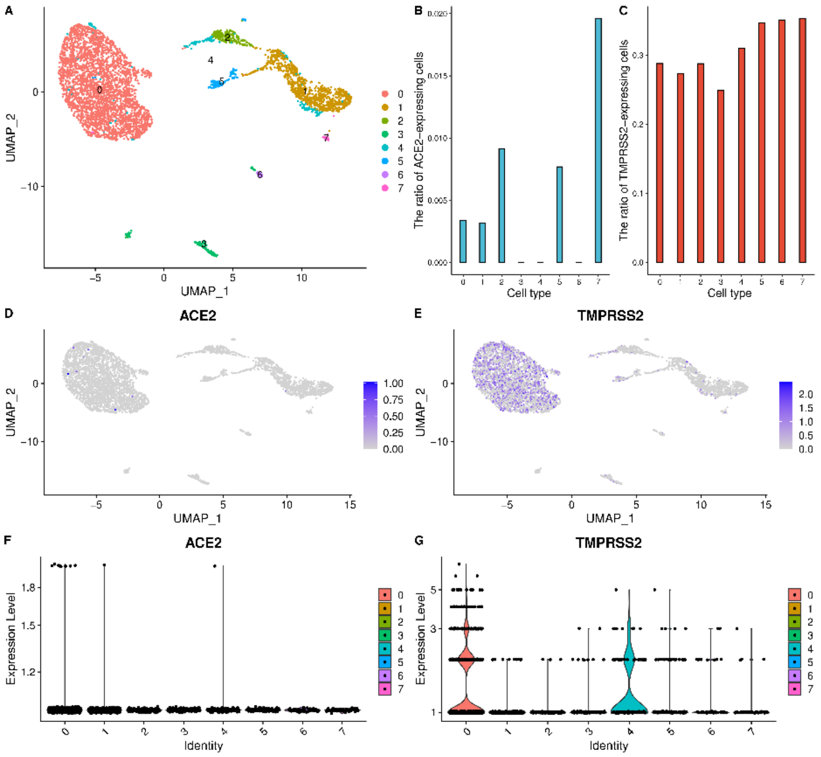


**Figure S7 The lung scRNA-seq data analysis results (donor 7). a**) UMAP visualization of clustering results for the lung cells. **b**) The ratio of ACE2-expressed cells in each cell cluster. **c**) The ratio of TMPRSS2-expressed cells in each cell cluster. **d**) ACE2 expression level in each cell cluster on the UMPA plot. **e**) TMPRSS2 expression level in each cell cluster on the UMPA plot. **f**) The expression distribution of ACE2 across each cell cluster. **g**) The expression distribution of TMPRSS2 across each cell cluster.

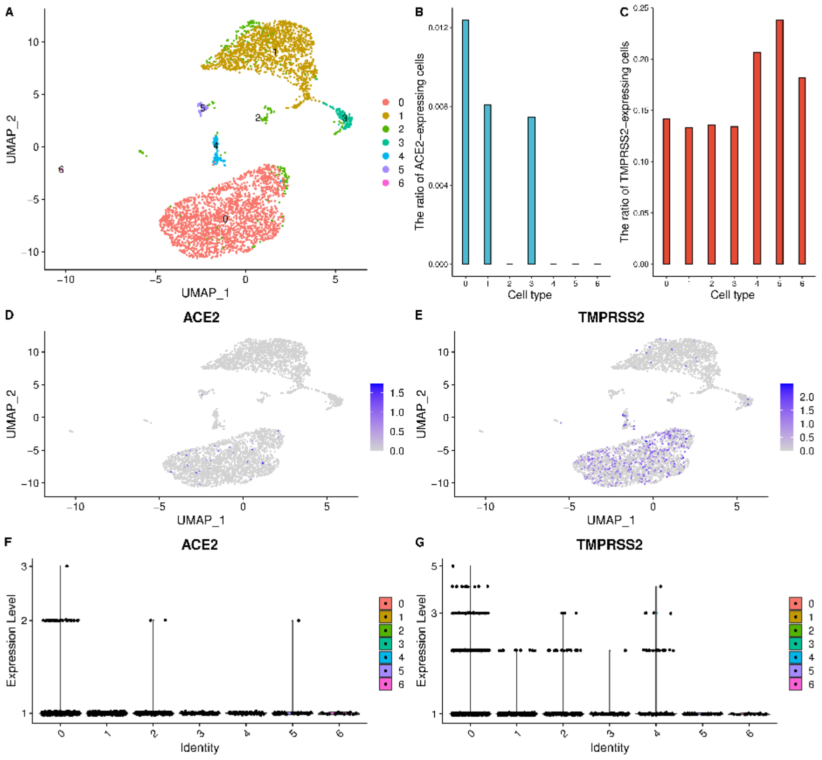


**Figure S8 The lung scRNA-seq data analysis results (donor 8). a**) UMAP visualization of clustering results for the lung cells. **b**) The ratio of ACE2-expressed cells in each cell cluster. **c**) The ratio of TMPRSS2-expressed cells in each cell cluster. **d**) ACE2 expression level in each cell cluster on the UMPA plot. **e**) TMPRSS2 expression level in each cell cluster on the UMPA plot. **f**) The expression distribution of ACE2 across each cell cluster. **g**) The expression distribution of TMPRSS2 across each cell cluster.

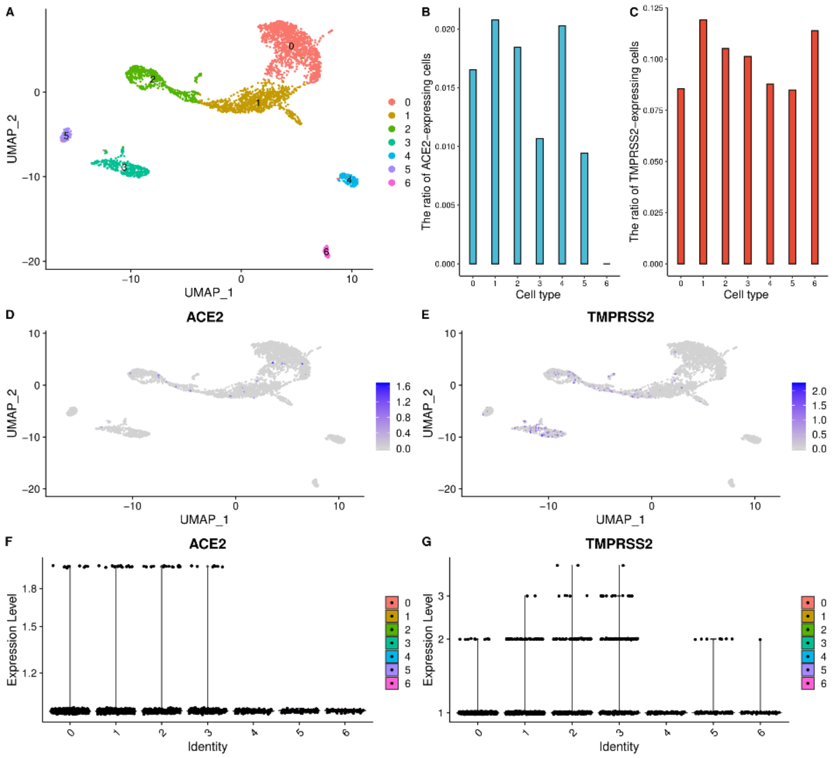


**Figure S9 High ACE2 and TMPRSS2 expression levels of the mesenchymal stromal cells, plasma cells in the nasal turbinate epithelial cells. a**) UMAP visualization of clustering results for the nasal turbinate epithelial cells. **b**) The ratio of ACE2-expressed cells in each cell cluster. **c**) The ratio of TMPRSS2-expressed cells in each cell cluster. **d**) ACE2 expression level in each cell cluster on the UMPA plot. **e**) TMPRSS2 expression level in each cell cluster on the UMPA plot. **f**) The expression distribution of ACE2 across each cell cluster. **g**) The expression distribution of TMPRSS2 across each cell cluster.

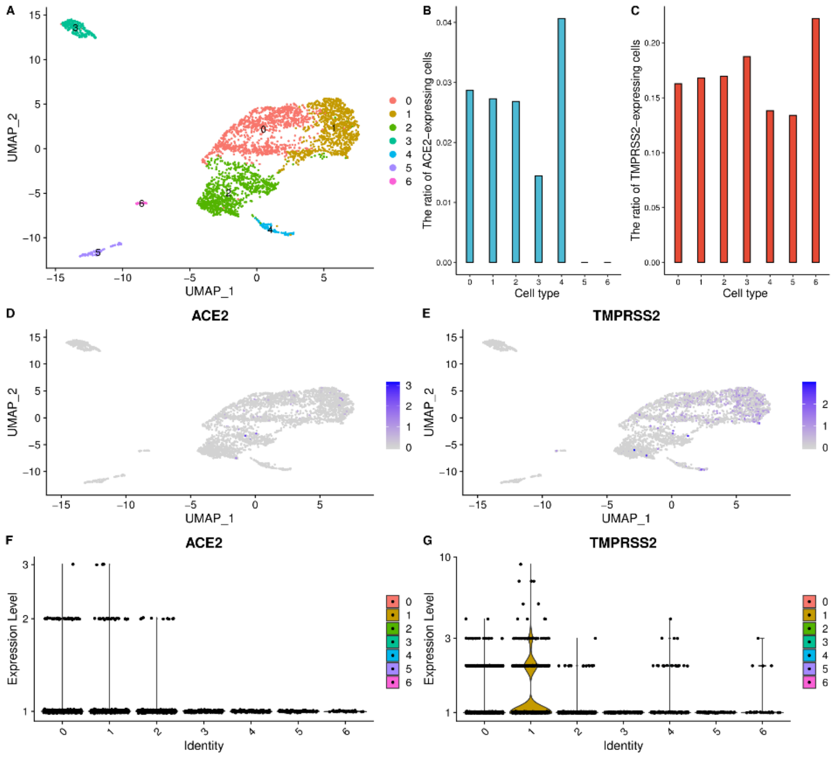


**Figure S10 High ACE2 and TMPRSS2 expression levels of the mesenchymal stromal cells, plasma cells in the nasal brushing epithelial cells.** **a**) UMAP visualization of clustering results for the nasal brushing epithelial cells. **b**) The ratio of ACE2-expressed cells in each cell cluster. **c**) The ratio of TMPRSS2-expressed cells in each cell cluster. **d**) ACE2 expression level in each cell cluster on the UMPA plot. **e**) TMPRSS2 expression level in each cell cluster on the UMPA plot. **f**) The expression distribution of ACE2 across each cell cluster. **g**) The expression distribution of TMPRSS2 across each cell cluster.

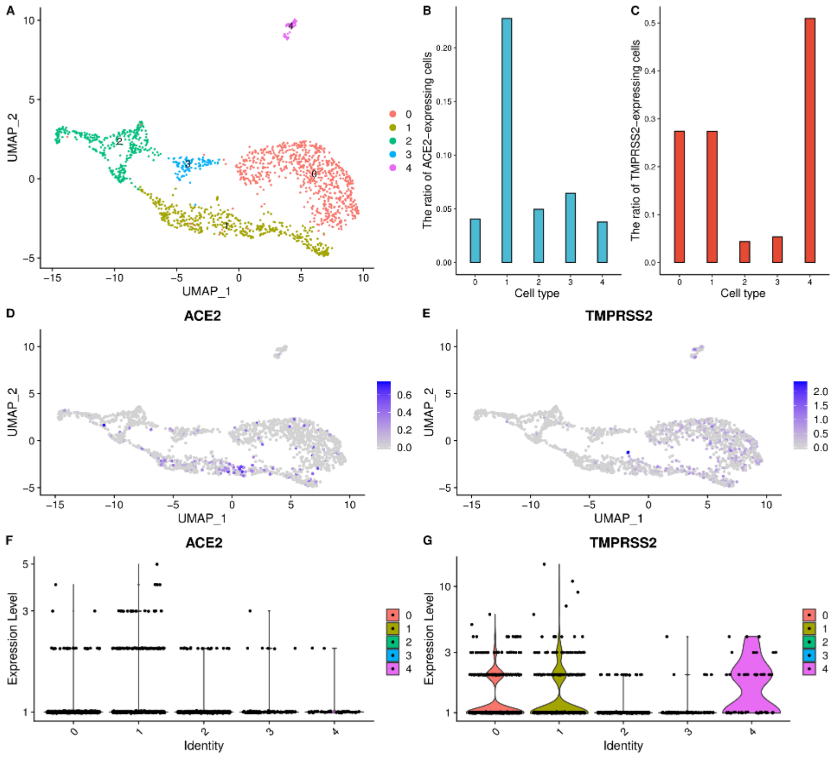


**Figure S11 High ACE2 and TMPRSS2 expression levels in the nasal airway epithelial cells. a)** UMAP visualization of clustering results for the airway epithelial cells. **b**) The ratio of ACE2-expressed cells in each cell cluster. **c**) The ratio of TMPRSS2-expressed cells in each cell cluster. **d**) ACE2 expression level in each cell cluster on the UMPA plot. **e**) TMPRSS2 expression level in each cell cluster on the UMPA plot. **f**) The expression distribution of ACE2 across each cell cluster. **g**) The expression distribution of TMPRSS2 across each cell cluster.

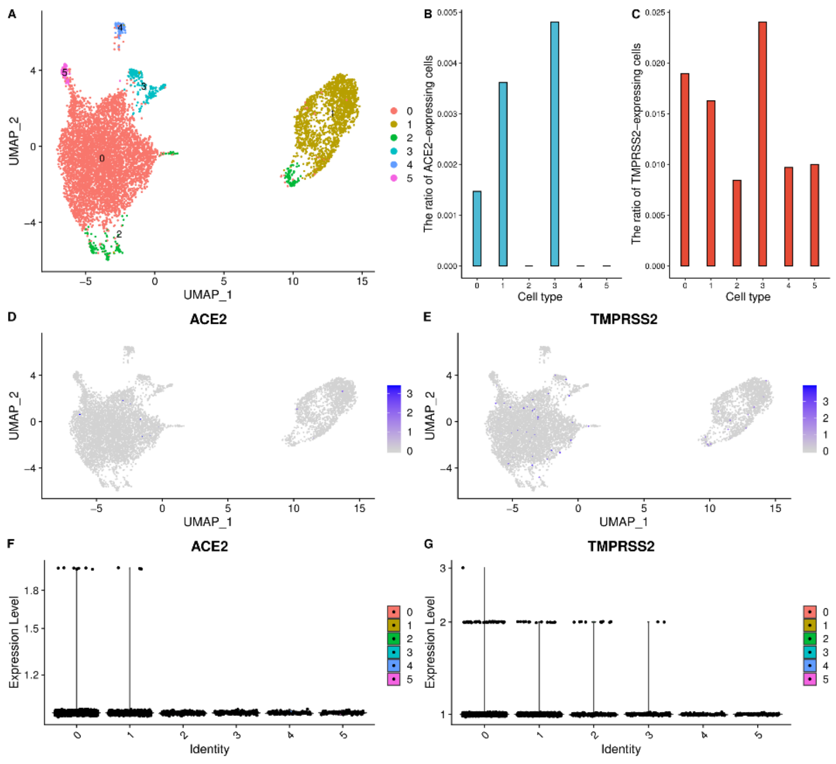


**Figure S12 The bronchus scRNA-seq data analysis results. a)** UMAP visualization of clustering results for the bronchus cells. **b**) The ratio of ACE2-expressed cells in each cell cluster. **c**) The ratio of TMPRSS2-expressed cells in each cell cluster. **d**) ACE2 expression level in each cell cluster on the UMPA plot. **e**) TMPRSS2 expression level in each cell cluster on the UMPA plot. **f**) The expression distribution of ACE2 across each cell cluster. **g**) The expression distribution of TMPRSS2 across each cell cluster.

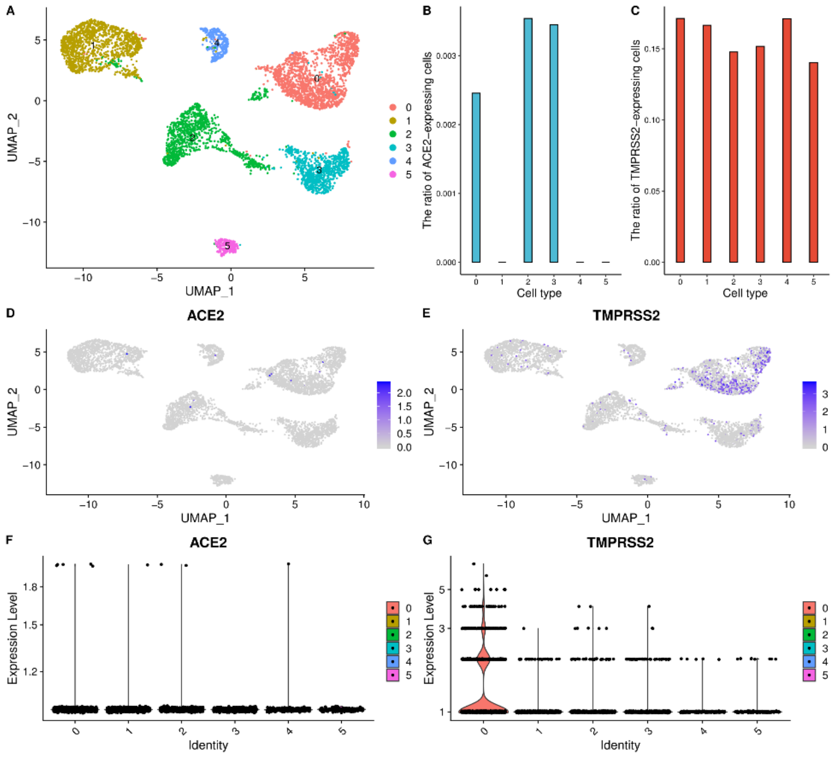


**Figure S13 The trachea scRNA-seq data analysis results. a**) UMAP visualization of clustering results for the trachea cells. **b**) The ratio of ACE2-expressed cells in each cell cluster. **c**) The ratio of TMPRSS2-expressed cells in each cell cluster. **d**) ACE2 expression level in each cell cluster on the UMPA plot. **e**) TMPRSS2 expression level in each cell cluster on the UMPA plot. **f**) The expression distribution of ACE2 across each cell cluster. **g**) The expression distribution of TMPRSS2 across each cell cluster.

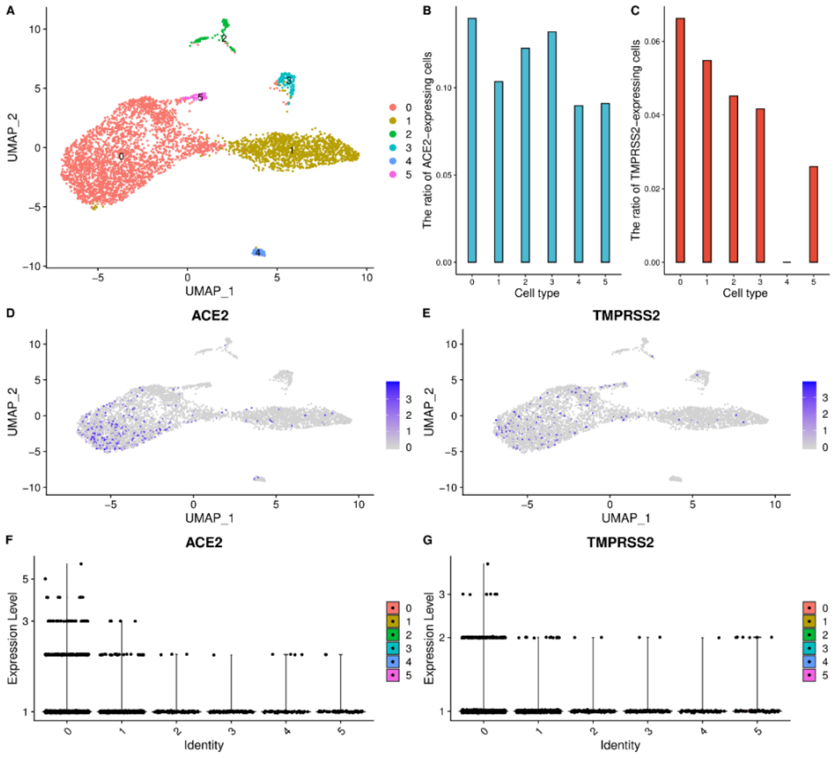


**Figure S14 High ACE2 and TMPRSS2 expression level of enterocyte progenitor cells and goblet cells in the jejunum.** a) UMAP visualization of clustering results for jejunum cells. **b**) The ratio of ACE2-expressed cells in each cell cluster. **c**) The ratio of TMPRSS2-expressed cells in each cell cluster. **d**) ACE2 expression level in each cell cluster on the UMPA plot. **e**) TMPRSS2 expression level in each cell cluster on the UMPA plot. **f**) The expression distribution of ACE2 across each cell cluster. **g**) The expression distribution of TMPRSS2 across each cell cluster.

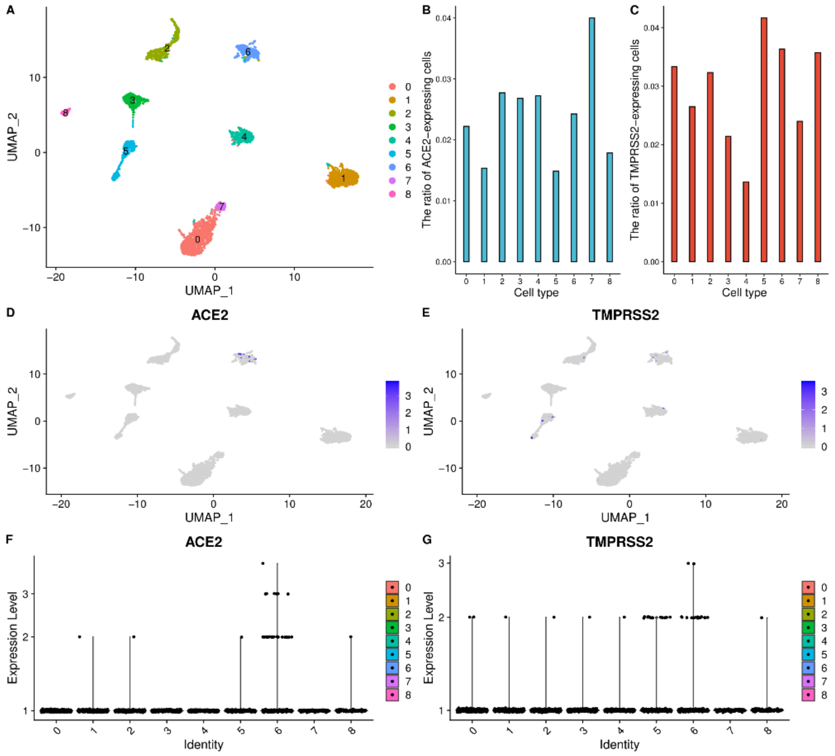


**Figure S15 High ACE2 and TMPRSS2 expression level of the intestinal epithelial stem cells and enterocyte progenitor cells in the ileum. a**) UMAP visualization of clustering results for ileum cells. **b**) The ratio of ACE2-expressed cells in each cell cluster. **c**) The ratio of TMPRSS2-expressed cells in each cell cluster. **d**) ACE2 expression level in each cell cluster on the UMPA plot. **e**) TMPRSS2 expression level in each cell cluster on the UMPA plot. **f**) The expression distribution of ACE2 across each cell cluster. **g**) The expression distribution of TMPRSS2 across each cell cluster.

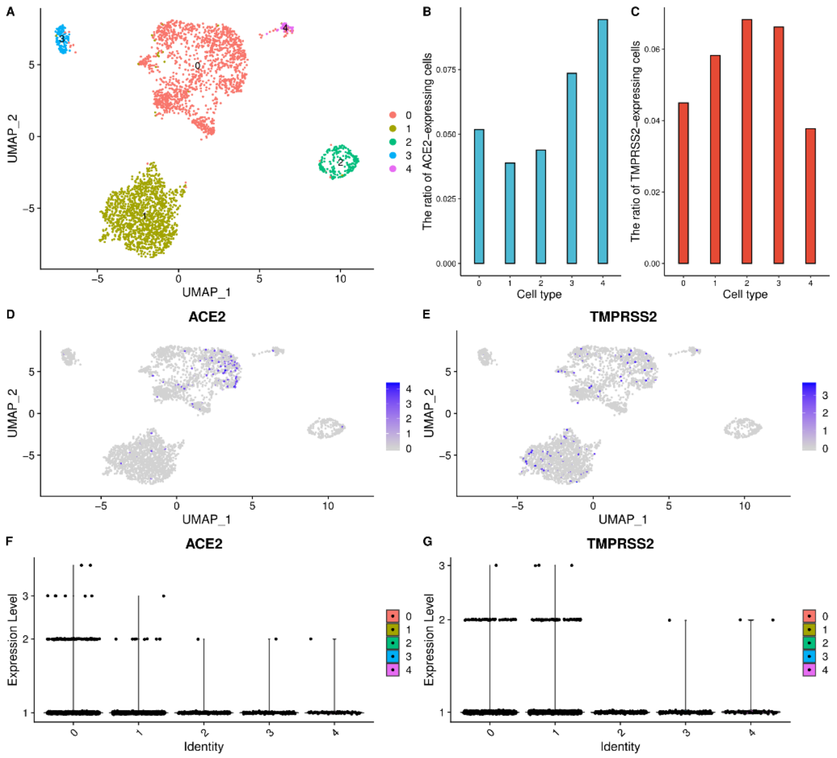


**Figure S16 High ACE2 and TMPRSS2 expression level of the intestinal LGR5+ stem cells, epithelial stem cells, enterocyte progenitor cells, tuft progenitor cells, and enteroendocrine cells in the duodenum.** **a**) UMAP visualization of clustering results for duodenum cells. **b**) The ratio of ACE2-expressed cells in each cell cluster. **c**) The ratio of TMPRSS2-expressed cells in each cell cluster. **d**) ACE2 expression level in each cell cluster on the UMPA plot. **e**) TMPRSS2 expression level in each cell cluster on the UMPA plot. **f**) The expression distribution of ACE2 across each cell cluster. **g**) The expression distribution of TMPRSS2 across each cell cluster.

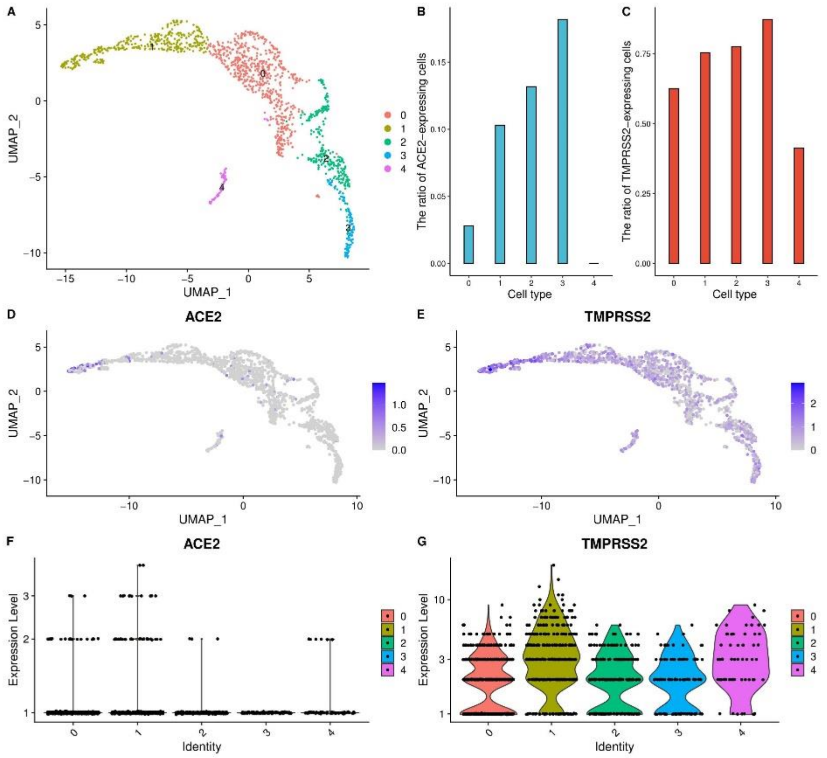


**Figure S17 High ACE2 and TMPRSS2 expression level of goblet progenitor cells, MKI67+ progenitor cells, enterocytes, and goblet cells in the rectum. a**) UMAP visualization of clustering results for rectum cells. **b**) The ratio of ACE2-expressed cells in each cell cluster. **c**) The ratio of TMPRSS2-expressed cells in each cell cluster. **d**) ACE2 expression level in each cell cluster on the UMPA plot. **e**) TMPRSS2 expression level in each cell cluster on the UMPA plot. **f**) The expression distribution of ACE2 across each cell cluster. **g**) The expression distribution of TMPRSS2 across each cell cluster.

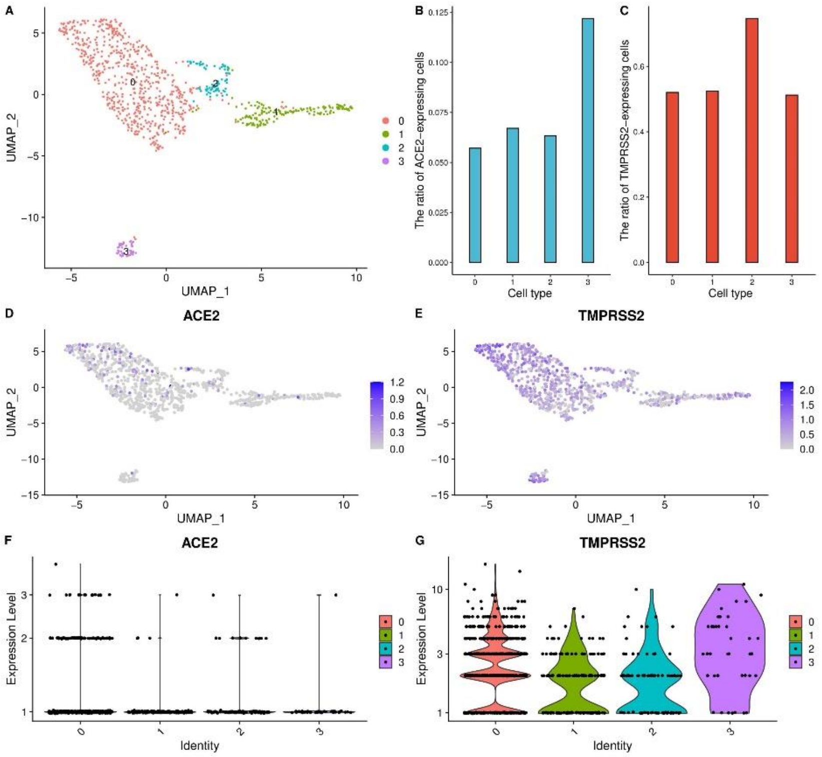


**Figure S18 High ACE2 and TMPRSS2 expression level of the enterocytes and goblet cells in the colon.** **a**) UMAP visualization of clustering results for colon cells. **b**) The ratio of ACE2-expressed cells in each cell cluster. **c**) The ratio of TMPRSS2-expressed cells in each cell cluster. **d**) ACE2 expression level in each cell cluster on the UMPA plot. **e**) TMPRSS2 expression level in each cell cluster on the UMPA plot. **f**) The expression distribution of ACE2 across each cell cluster. **g**) The expression distribution of TMPRSS2 across each cell cluster.

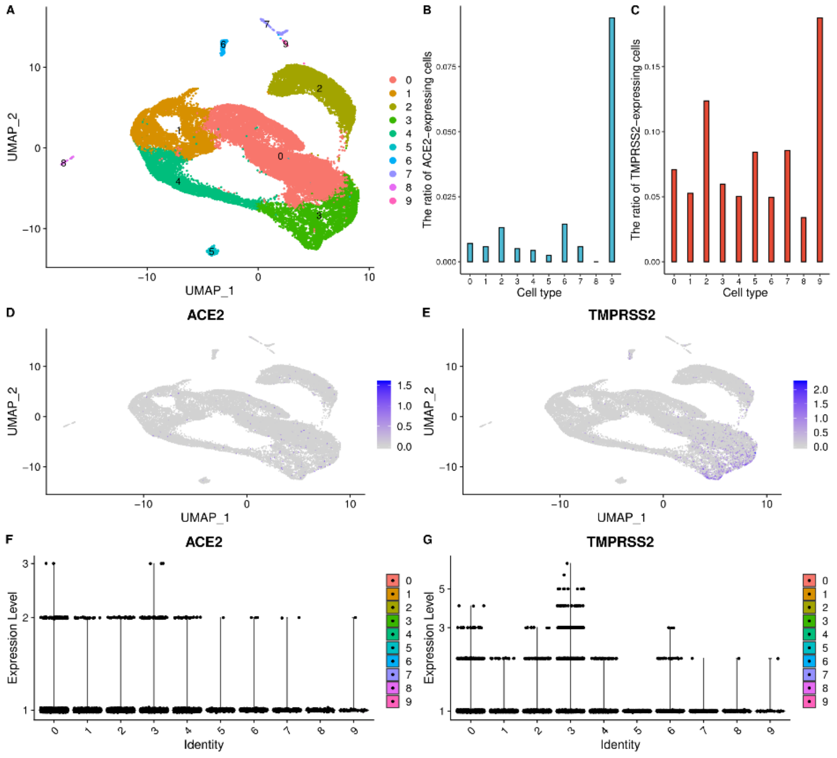


**Figure S19 High ACE2 and TMPRSS2 expression level of the secretory progenitor cells in the esophagus.** **a**) UMAP visualization of clustering results for esophagus cells. **b**) The ratio of ACE2-expressed cells in each cell cluster. **c**) The ratio of TMPRSS2-expressed cells in each cell cluster. **d**) ACE2 expression level in each cell cluster on the UMPA plot. **e**) TMPRSS2 expression level in each cell cluster on the UMPA plot. **f**) The expression distribution of ACE2 across each cell cluster. **g**) The expression distribution of TMPRSS2 across each cell cluster.

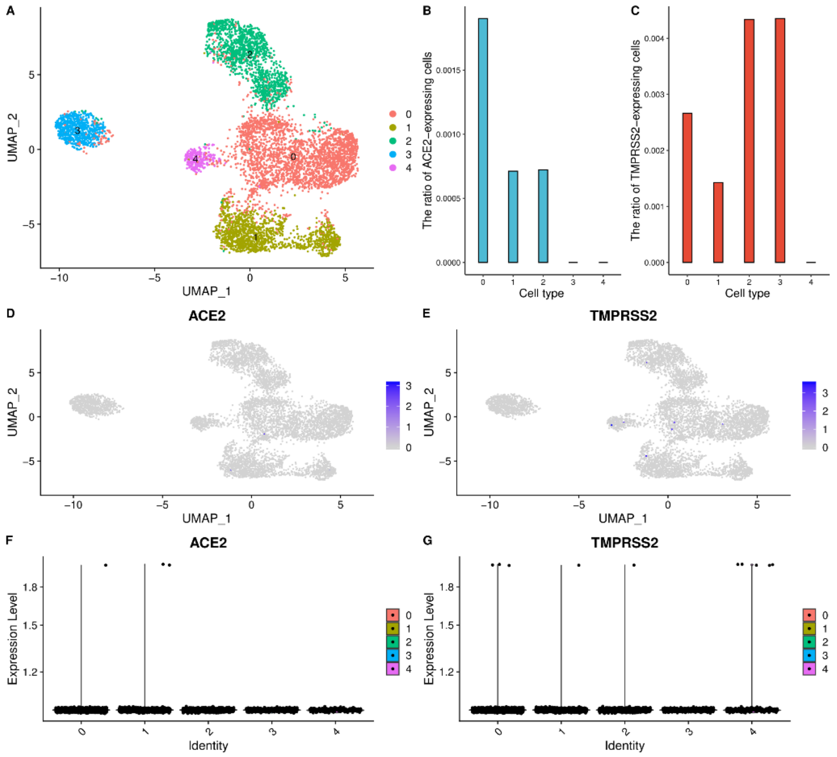


**Figure S20 The liver scRNA-seq data analysis results. a**) UMAP visualization of clustering results for liver cells. **b**) The ratio of ACE2-expressed cells in each cell cluster. **c**) The ratio of TMPRSS2-expressed cells in each cell cluster. **d**) ACE2 expression level in each cell cluster on the UMPA plot. **e**) TMPRSS2 expression level in each cell cluster on the UMPA plot. **f**) The expression distribution of ACE2 across each cell cluster. **g**) The expression distribution of TMPRSS2 across each cell cluster.

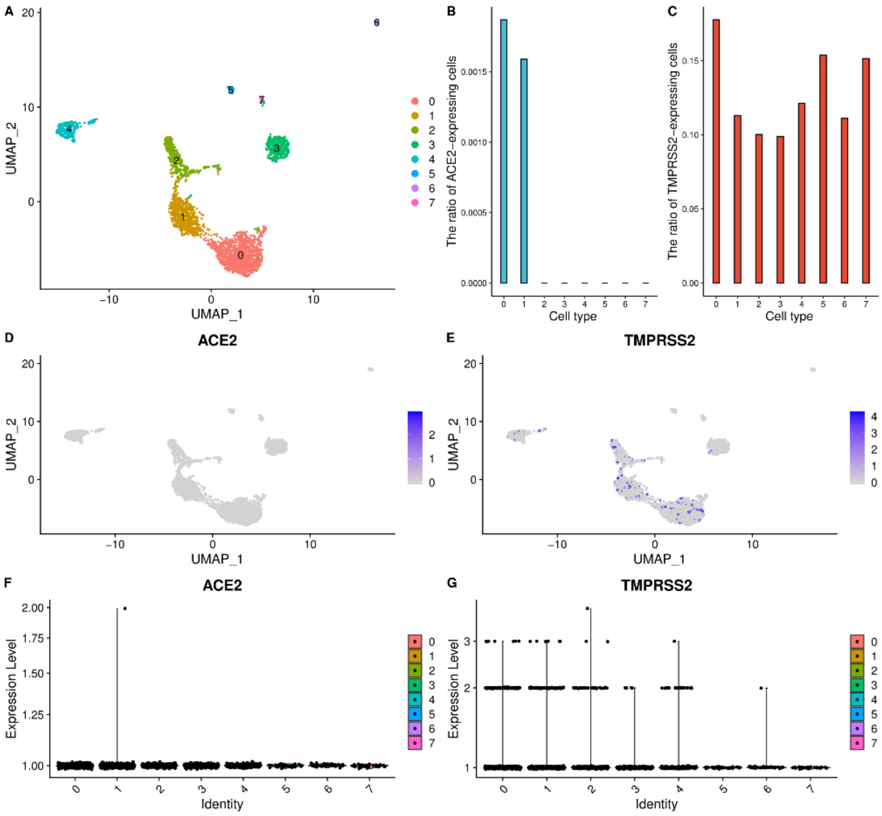

**Figure S21 The stomach scRNA-seq data analysis results. a**) UMAP visualization of clustering results for stomach cells. **b**) The ratio of ACE2-expressed cells in each cell cluster. **c**) The ratio of TMPRSS2-expressed cells in each cell cluster. **d**) ACE2 expression level in each cell cluster on the UMPA plot. **e**) TMPRSS2 expression level in each cell cluster on the UMPA plot. **f**) The expression distribution of ACE2 across each cell cluster. **g**) The expression distribution of TMPRSS2 across each cell cluster.

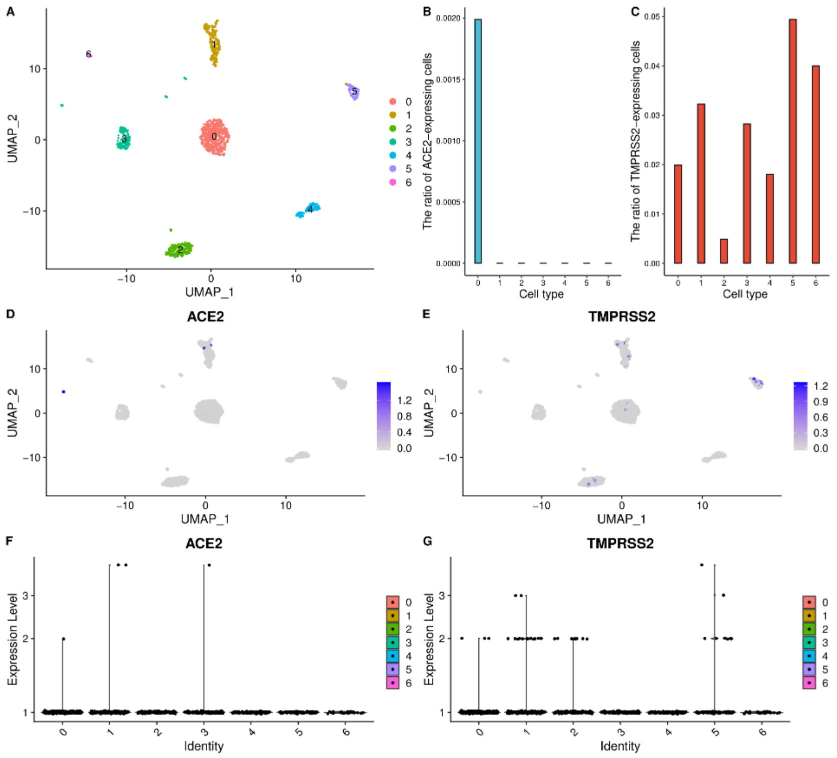


**Figure S22 The pancreatic islets scRNA-seq data analysis results.** **a**) UMAP visualization of clustering results for pancreatic islets cells. **b**) The ratio of ACE2-expressed cells in each cell cluster. **c**) The ratio of TMPRSS2-expressed cells in each cell cluster. **d**) ACE2 expression level in each cell cluster on the UMPA plot. **e**) TMPRSS2 expression level in each cell cluster on the UMPA plot. **f**) The expression distribution of ACE2 across each cell cluster. **g**) The expression distribution of TMPRSS2 across each cell cluster.

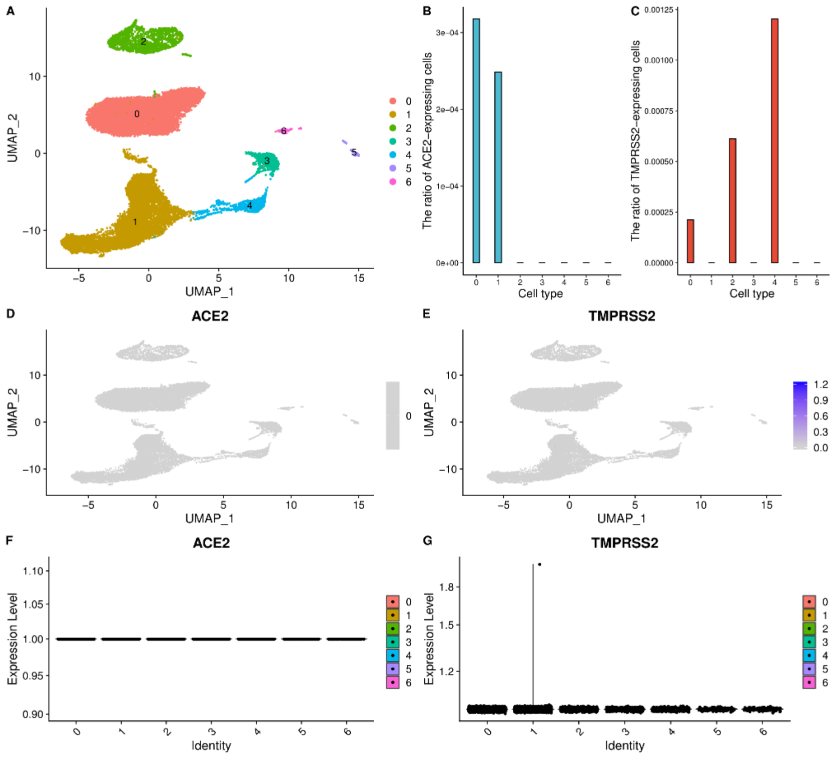


**Figure S23 The hippocampus scRNA-seq data analysis results. a**) UMAP visualization of clustering results for hippocampus cells. **b**) The ratio of ACE2-expressed cells in each cell cluster. **c**) The ratio of TMPRSS2-expressed cells in each cell cluster. **d**) ACE2 expression level in each cell cluster on the UMPA plot. **e**) TMPRSS2 expression level in each cell cluster on the UMPA plot. **f**) The expression distribution of ACE2 across each cell cluster. **g**) The expression distribution of TMPRSS2 across each cell cluster.

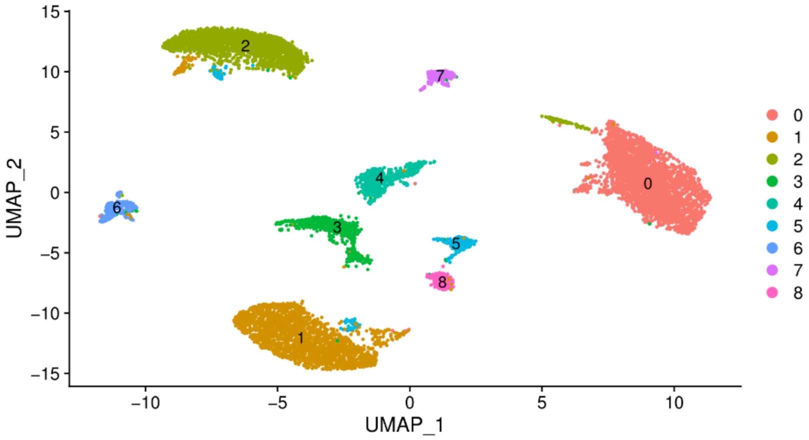


**Figure S24 UMAP visualization of clustering results for the cerebellum cells.**

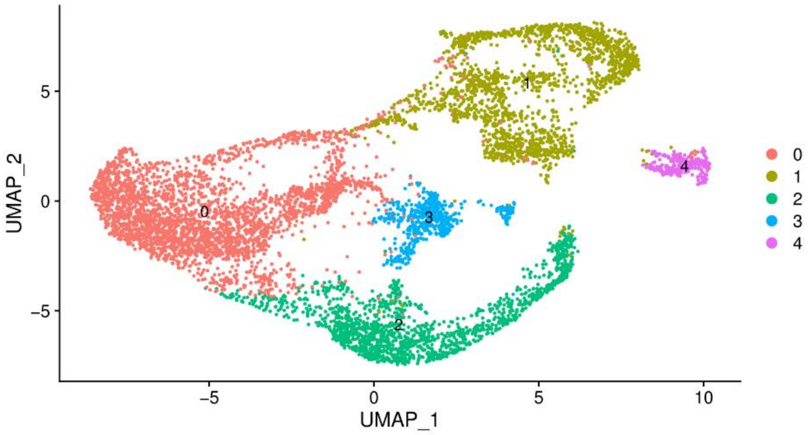


**Figure S25 UMAP visualization of clustering results for the spinal cord cells.**

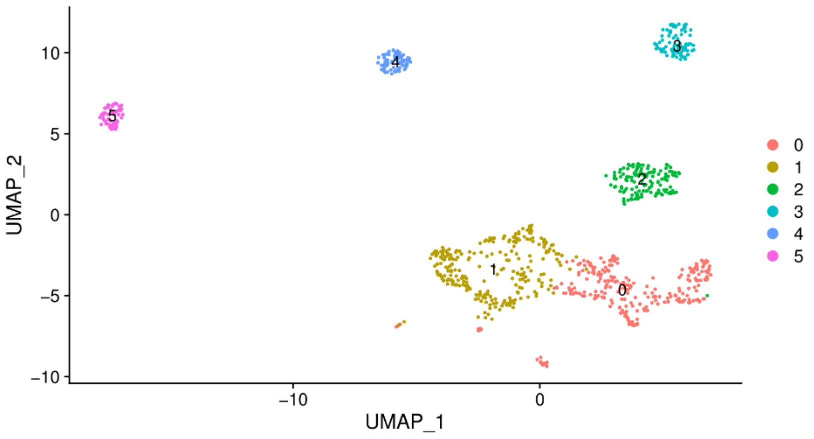


**Figure S26 UMAP visualization of clustering results for the neuronal epithelium cells.**

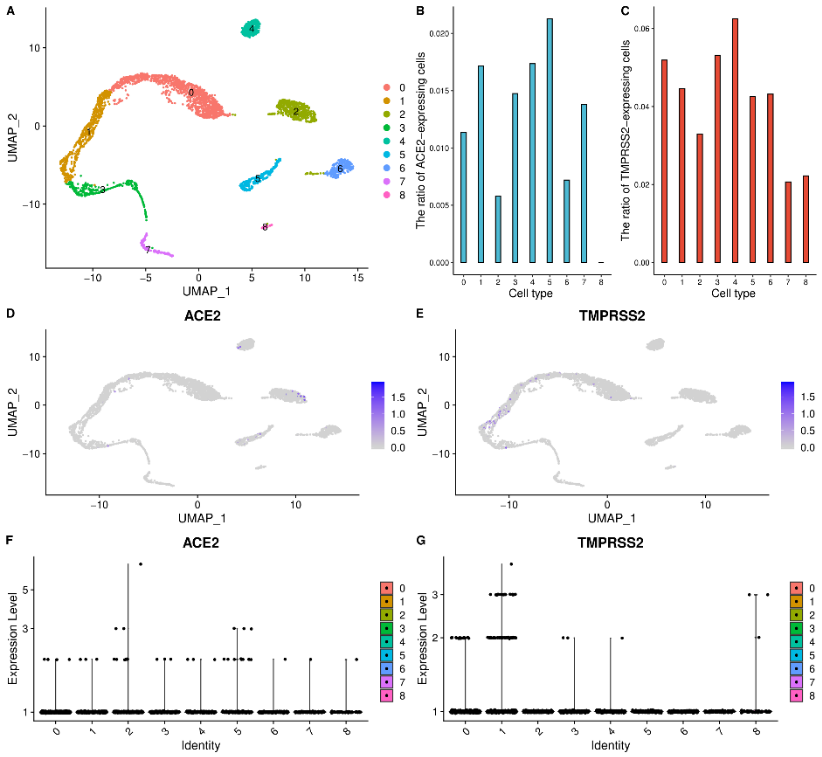


**Figure S27 High ACE2 and TMPRSS2 expression level of spermatogonium, peritubular myoid cells, testis somatic cells, and spermatogonial stem cells in the testis. a**) UMAP visualization of clustering results for testis cells. **b**) The ratio of ACE2-expressed cells in each cell cluster. **c**) The ratio of TMPRSS2-expressed cells in each cell cluster. **d**) ACE2 expression level in each cell cluster on the UMPA plot. **e**) TMPRSS2 expression level in each cell cluster on the UMPA plot. **f**) The expression distribution of ACE2 across each cell cluster. **g**) The expression distribution of TMPRSS2 across each cell cluster.

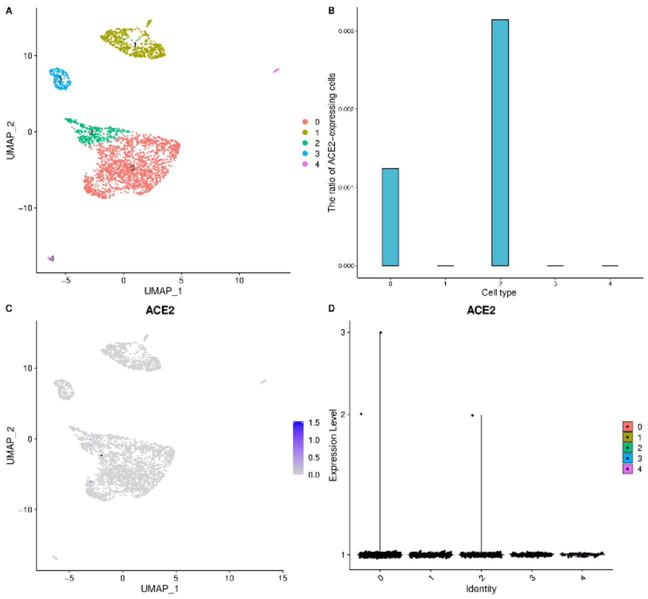


**Figure S28 The ovary scRNA-seq data analysis results. a**) UMAP visualization of clustering results for ovary cells. **b**) The ratio of ACE2-expressed cells in each cell cluster. **c**) ACE2 expression level in each cell cluster on the UMPA plot. **d**) The expression distribution of ACE2 across each cell cluster.

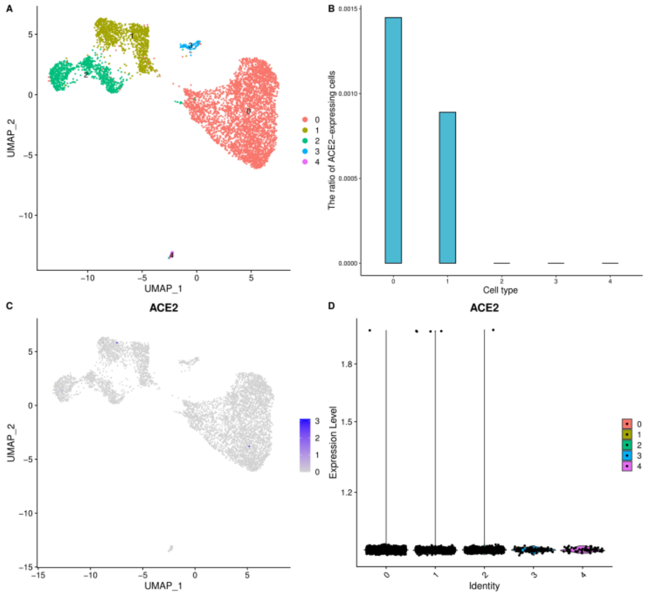


**Figure S29 The uterus scRNA-seq data analysis results. a**) UMAP visualization of clustering results for uterus cells. **b**) The ratio of ACE2-expressed cells in each cell cluster. **c**) ACE2 expression level in each cell cluster on the UMPA plot. **d**) The expression distribution of ACE2 across each cell cluster.

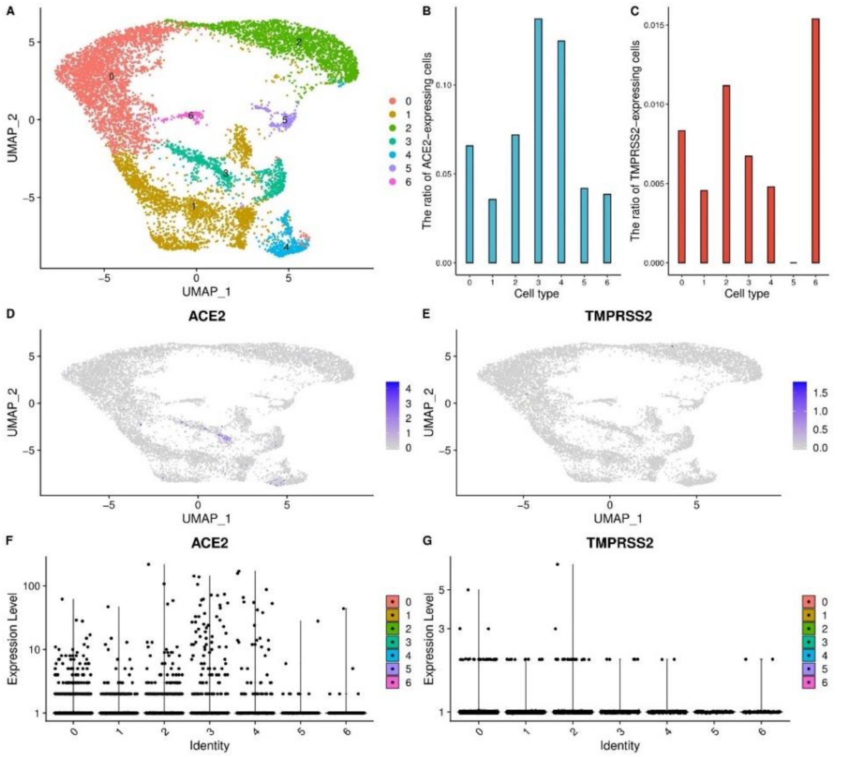


**Figure S30 High ACE2 and TMPRSS2 expression level of the cardiomyocytes and cardiovascular progenitor cells in the heart. a**) UMAP visualization of clustering results for the heart cells. **b**) The ratio of ACE2-expressed cells in each cell cluster. **c**) The ratio of TMPRSS2-expressed cells in each cell cluster. **d**) ACE2 expression level in each cell cluster on the UMPA plot. **e**) TMPRSS2 expression level in each cell cluster on the UMPA plot. **f**) The expression distribution of ACE2 across each cell cluster. **g**) The expression distribution of TMPRSS2 across each cell cluster.

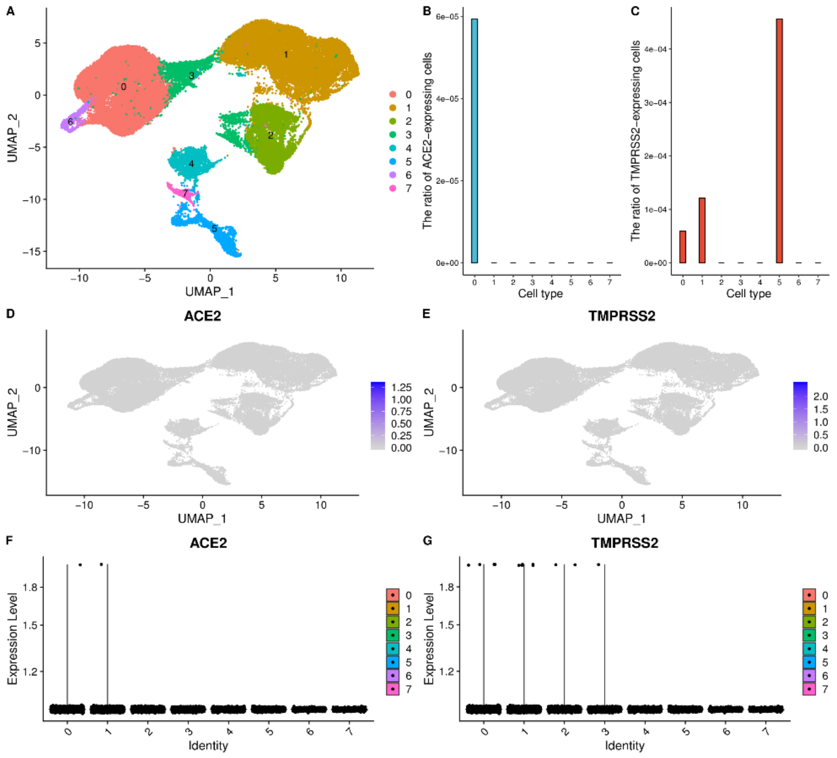


**Figure S31 The spleen scRNA-seq data analysis results. a**) UMAP visualization of clustering results for the spleen cells. **b**) The ratio of ACE2-expressed cells in each cell cluster. **c**) The ratio of TMPRSS2-expressed cells in each cell cluster. **d**) ACE2 expression level in each cell cluster on the UMPA plot. **e**) TMPRSS2 expression level in each cell cluster on the UMPA plot. **f**) The expression distribution of ACE2 across each cell cluster. **g**) The expression distribution of TMPRSS2 across each cell cluster.

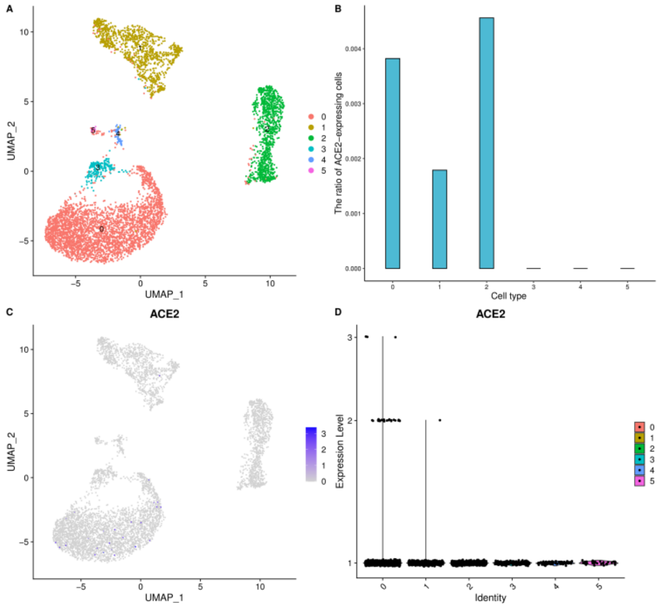


**Figure S32 The artery scRNA-seq data analysis results. a**) UMAP visualization of clustering results for the spleen cells. **b**) The ratio of ACE2-expressed cells in each cell cluster. **c**) ACE2 expression level in each cell cluster on the UMPA plot. **d**) The expression distribution of ACE2 across each cell cluster.

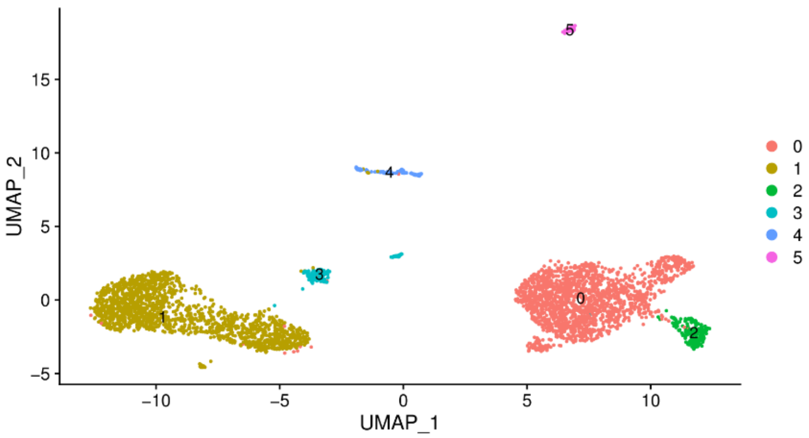


**Figure S33 UMAP visualization of clustering results for the peripheral blood cells.**

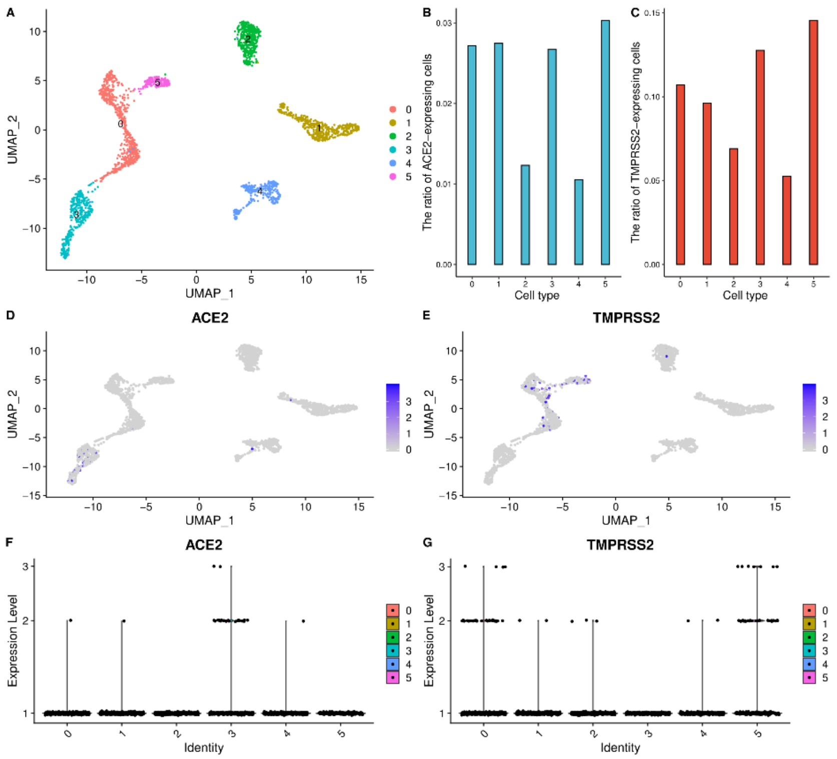


**Figure S34 High ACE2 and TMPRSS2 expression level of the nephron epithelial cells, epithelial cells, endothelial cells, and mesangial cells in the kidney. a**) UMAP visualization of clustering results for the kidney cells. **b**) The ratio of ACE2-expressed cells in each cell cluster. **c**) The ratio of TMPRSS2-expressed cells in each cell cluster. **d**) ACE2 expression level in each cell cluster on the UMPA plot. **e**) TMPRSS2 expression level in each cell cluster on the UMPA plot. **f**) The expression distribution of ACE2 across each cell cluster. **g**) The expression distribution of TMPRSS2 across each cell cluster.

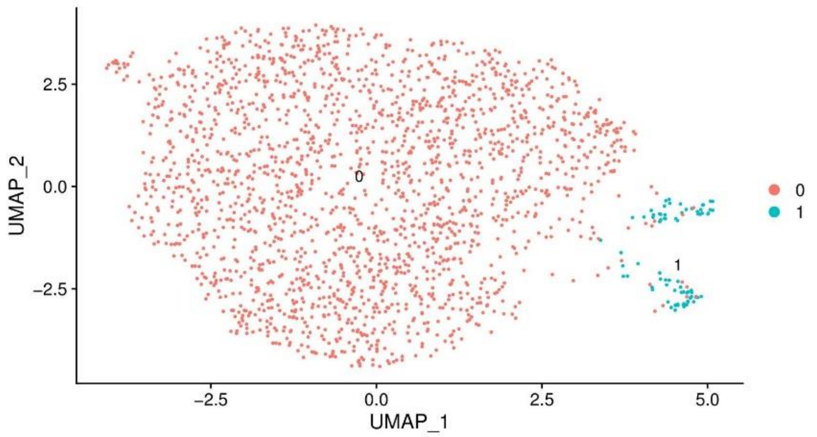


**Figure S35 UMAP visualization of clustering results for the ureter cells.**

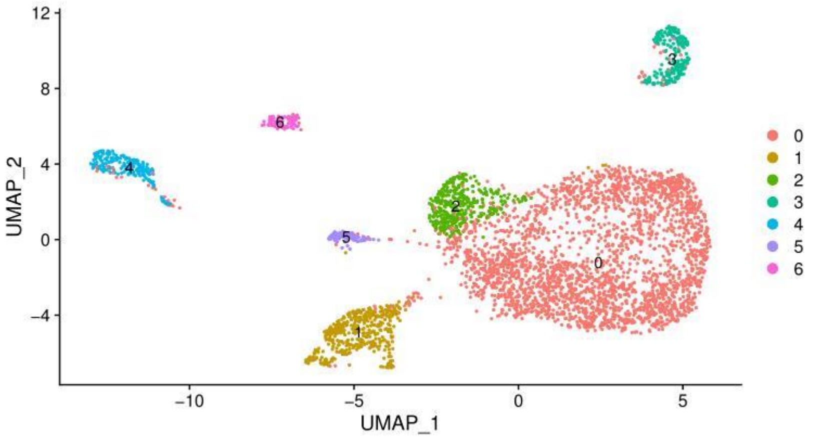


**Figure S36 UMAP visualization of clustering results for the prostate cells.**

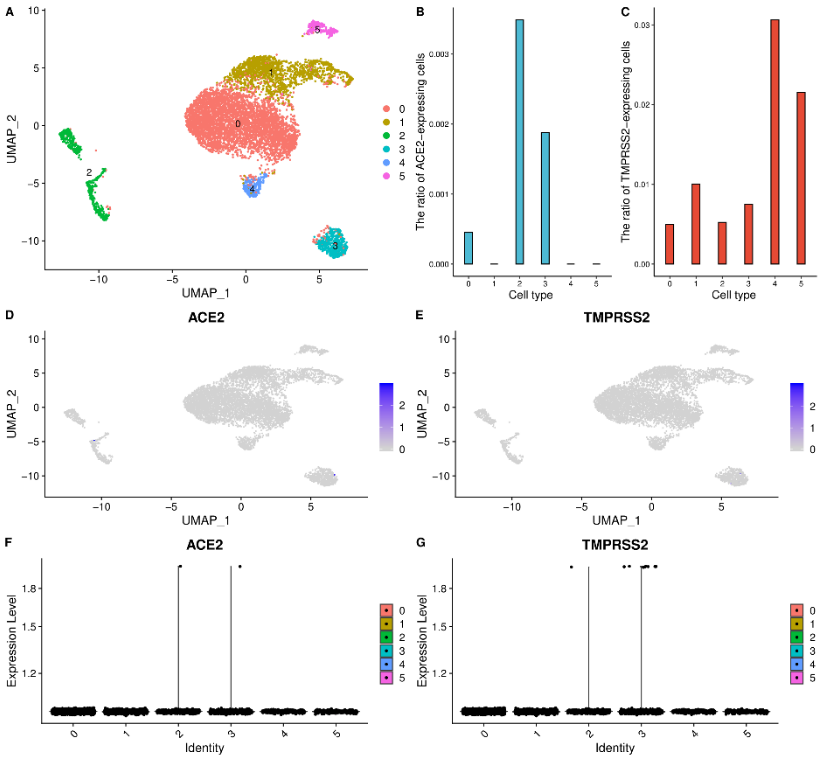


**Figure S37 The thyroid gland scRNA-seq data analysis results. a**) UMAP visualization of clustering results for the thyroid gland cells. **b**) The ratio of ACE2-expressed cells in each cell cluster. **c**) The ratio of TMPRSS2-expressed cells in each cell cluster. **d**) ACE2 expression level in each cell cluster on the UMPA plot. **e**) TMPRSS2 expression level in each cell cluster on the UMPA plot. **f**) The expression distribution of ACE2 across each cell cluster. **g**) The expression distribution of TMPRSS2 across each cell cluster.

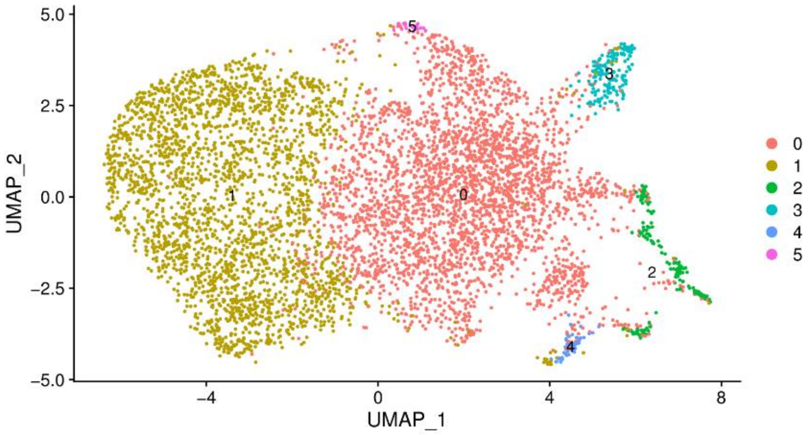


**Figure S38 UMAP visualization of clustering results for the thymus gland cells.**

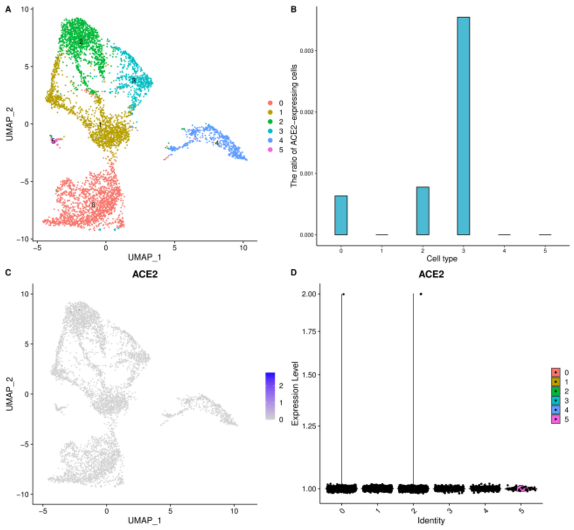


**Figure S39 The muscle scRNA-seq data analysis results. a**) UMAP visualization of clustering results for the muscle cells. **b**) The ratio of ACE2-expressed cells in each cell cluster. **c**) ACE2 expression level in each cell cluster on the UMPA plot. **d**) The expression distribution of ACE2 across each cell cluster.

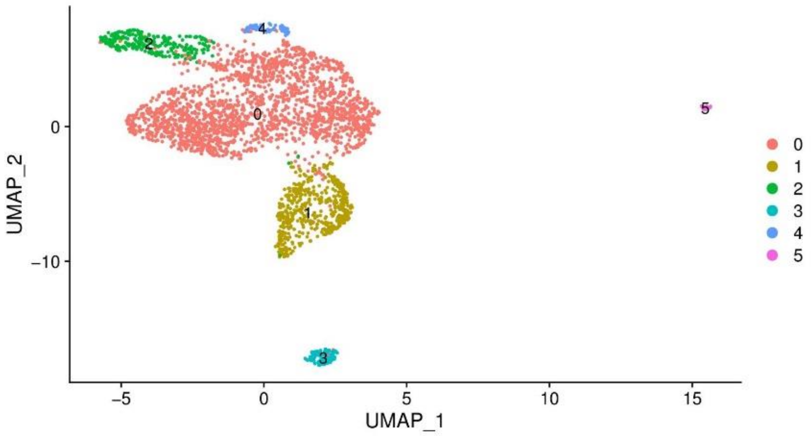


**Fig. S40 UMAP visualization of clustering results for the lymph nodes cells.**

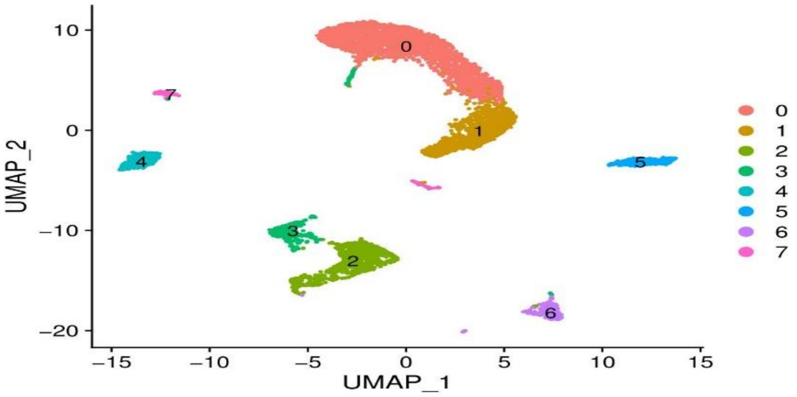


**Figure S41 UMAP visualization of clustering results for the tonsil dendritic cells.**

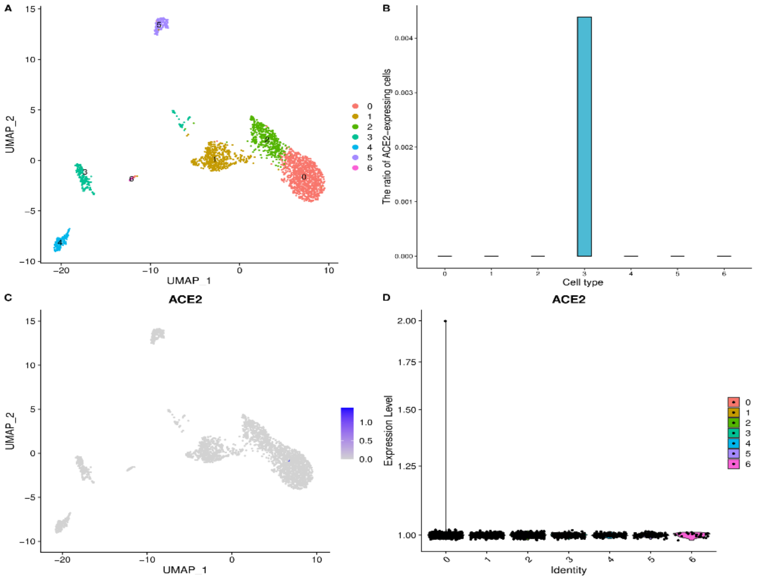


**Figure S42 The bone marrow scRNA-seq data analysis results. a**) UMAP visualization of clustering results for the bone marrow cells. **b**) The ratio of ACE2-expressed cells in each cell cluster. **c**) ACE2 expression level in each cell cluster on the UMPA plot. **d**) The expression distribution of ACE2 across each cell cluster.
